# TORCphysics: A physical model of DNA-topology-controlled gene expression

**DOI:** 10.1101/2025.06.04.656888

**Authors:** Victor Velasco-Berrelleza, Penn Faulkner Rainford, Aalap Mogre, Craig J. Benham, Charles J. Dorman, Carsten Kröger, Susan Stepney, Sarah A. Harris

## Abstract

DNA superhelicity and transcription are intimately related because changes to DNA topology can influence gene expression and vice versa. Information is transferred through the modulation of local DNA torsional stress, where the expression of one gene may influence the superhelical level of neighbouring genes, either promoting or repressing their expression. In this work, we introduce a one-dimensional physical model that simulates supercoiling-mediated regulation. This TORCphysics model takes as input a genome architecture represented either by a plasmid or by a linear DNA sequence with ends constrained under specific biological conditions, and computes the molecule’s output. Our findings demonstrate that the expression profiles of genes are directly influenced by the gene circuit design, including gene location, the positions of topological barriers, promoter sequences, and topoisomerase activity. The novelty that TORCphysics offers is versatility, where users can define distinct activity models for different types of proteins and protein binding sites. The aim of this research is to establish a flexible framework for developing physical simulations of gene circuits to deepen our comprehension of the intricate mechanisms involved in gene regulation.

## 1 Introduction

The level of over- or underwinding of the DNA double helix, known as supercoiling, is a fundamental topological property of DNA. In bacteria, DNA is typically maintained in a negatively supercoiled (underwound) state. The DNA is therefore under torsional stress, as turns have been removed from the double helix [1]. This underwound state is regulated by a delicate balance between the activities of *topoisomerases*, enzymes that modulate superhelical stress. In bacteria, topoisomerase I (type IA) removes supercoils via single-strand breaks, while gyrase (type IIA) introduces negative supercoils by cutting and rejoining both strands in an ATP-dependent manner [2–5]. In the absence of ATP, gyrase is capable of relaxing negatively supercoiled DNA [6]. A recent FRET-based kinetics study on fluorescent labeled plasmids shows that both enzymes follow classical Michaelis-Menten kinetics in *Escherichia coli* [7].

DNA supercoiling is intimately related to transcription. It can control promoter activity and influence various stages of transcription [4, 8, 9]. During closed transcription complex formation, DNA supercoiling influences the binding rate of RNA polymerases (RNAPs) by altering the geometric orientation of the −10 and −35 promoter elements that are separated by the promoter spacer length [10]. In open-complex formation, negative supercoiling may reduce the free energy required for strand separation at the promoter, thereby facilitating transcription bubble formation [11–13]. During elongation, RNAP acts as a mobile topological barrier, generating positive supercoils downstream and negative supercoils upstream, which can propagate and influence the expression of distal genes as well as the gene being transcribed [14–18]. Positive supercoils can inhibit transcription and stall RNAPs, while topoisomerases help alleviate torsional stress, thereby enabling transcription to resume [14, 15, 19–21]. Recent evidence suggests that topoisomerase I physically interacts with transcribing RNAPs upstream, and this interaction may play a crucial role in transcript elongation [22–25].

Quantitatively predicting how DNA supercoiling modulates gene expression introduces further complexity into models that rely solely on transcription factors to promote or repress gene expression [26]. Several models have emerged to investigate the coupling between supercoiling and transcription [14, 27–36]. These models vary in focus: some emphasize transcription initiation [14, 27, 28, 34], some explore the collective motion of RNAPs during elongation and its effects on transcription bursting [30–32, 34], some investigate how transcription is influenced by supercoiling diffusion [29, 30, 36].

In this work, we introduce TORCphysics, a novel, versatile, and fast one-dimensional physical model that accounts for the interactions between DNA and proteins such as RNAP and topoisomerases through DNA supercoiling, with a particular focus on transcription. Our model incorporates the stochastic binding of topoisomerases on DNA, the interaction between topoisomerase I and transcribing RNAPs, and the impact of DNA supercoiling during transcription initiation and elongation. We assume all changes to DNA superhelicity introduced by bound proteins are expressed exclusively as changes in twist: we do not account for changes in writhe or any resulting structures such as plectonemes. Even with these assumptions, our results provide mechanistic insight into gene regulation through DNA supercoiling. By revisiting previous experimental studies, we calibrate our model and propose potential mechanisms that may explain observed experimental phenomena. Our results demonstrate that DNA-binding proteins both contribute and respond to local DNA supercoiling, revealing complex mechanisms even in simple genetic architectures such as single-gene systems. TORCphysics provides a flexible platform for simulating genetic circuits, enabling users to adjust the complexity of molecular interactions to investigate, compare, and explain experimental findings.

## 2 Methods – CoSMoS Overview

To develop the TORCphysics system, we follow the CoSMoS (Complex Systems Modelling and Simulation) approach, as described in detail in [37]. Here, we provide a brief overview of the approach.

CoSMoS involves a variety of components: domain, domain model, platform model, simulation platform, and results model (Figure 1). Each of these components plays a particular role the design, implementation, and use of a simulation.

**Figure 1.**
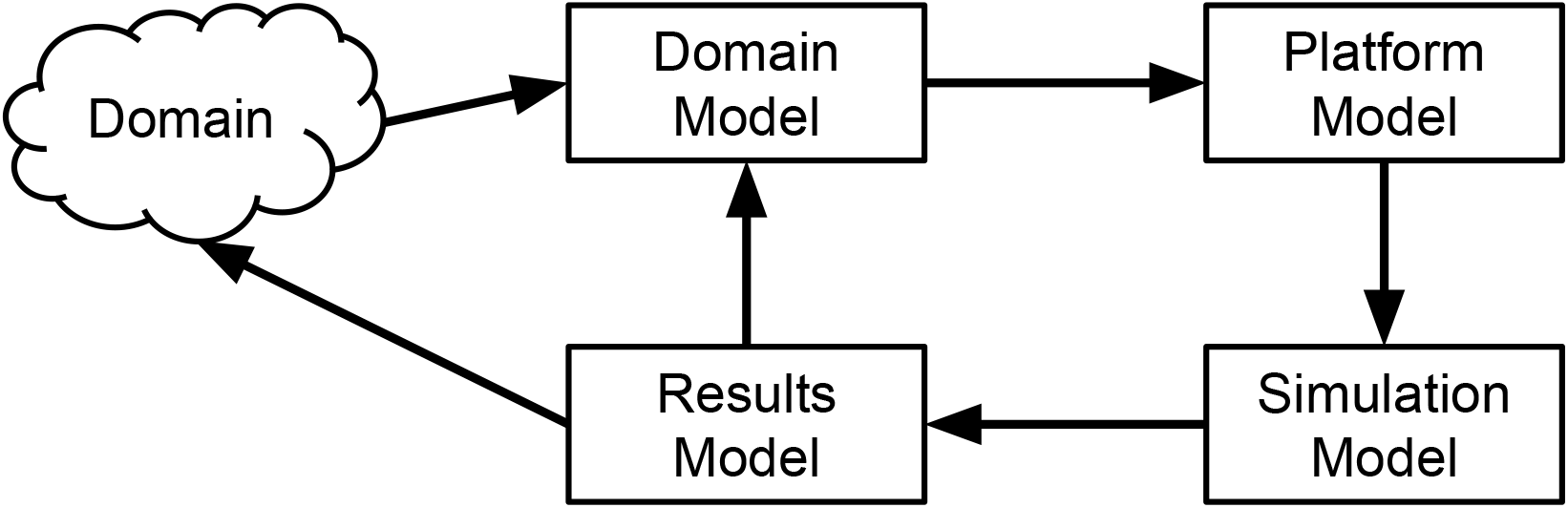
The modelling components of a CoSMoS simulation development. The CoSMoS approach formalizes the construction of models and simulations of complex systems and here is applied to transcription regulation through DNA supercoiling in the TORCphysics scheme.

### Domain

This is (a view of) the particular real world system being simulated. The TORCphysics domain is DNA supercoiling and transcription in bacteria. It is described in section 1.

### Domain Model

This is a scientific model of the relevant parts of the domain. It can incorporate understanding gained from domain experiments and observations, and hypotheses of the particular underlying mechanisms of interest. Crucially, the domain model is expressed in terms of relevance to the domain scientists, is validated by them as an accurate representation of the domain, and does not include simulation implementation details.

The TORCphysics domain model is a model of the physical processes involved in supercoiling and gene expression, given in terms of Ordinary Differential Equations. It is detailed in section 3.1.

### Platform Model

This is a software engineering model, or requirements specification, of the simulation software to be developed. It maps the domain model into computational terms, and includes necessary implementation detail. The mapping may make assumptions, approximations, and other changes, in order to make a computational simulation feasible. For example, a continuous time domain model will need to be converted to a discrete time platform model.

The platform model captures the low level (possibly hypothesised) domain model mechanisms, explicitly omitting the high level (emergent) properties observed in domain experiments. This ensures that those emergent properties are not ‘hardcoded’ into the simulation, thereby allowing them to emerge (or not) from the hypothesised mechanisms. For example, the domain model might include gene expression levels, hypothesised to be caused by particular supercoiling mechanisms. The platform model would include the mechanisms, but not hardcode the expression levels: these would emerge as consequences of the mechanisms during simulation experiment runs. If they match the domain values, this provides evidence to support the mechanisms; if they do not, other mechanisms may need to be sought.

The TORCPhysics platform model is described in section 3.2.

### Simulation Platform

This is the software implementation of the platform model, calibrated against experimental data as required. The calibrated platform is used to perform simulation experiments. Note the logical distance of the simulation code from the domain: it is not a direct ‘coding up’ of domain concepts. A CoSMoS simulator is carefully engineered in a manner explicitly designed to ensure that any emergent properties in the results have not been accidentally hardcoded in, hence the hypothesised mechanisms can be rigorously evaluated.

The TORCphysics simulation platform code is available from https://github.com/Victor-93/TORCphysics/tree/TORCphysics_paper. The TORCPhysics simulation platform’s calibration is detailed in section 3.3. More information about the TORCphysics repository and code availability is provided in Supplementary Section 2.

### Results Model

This descriptive model captures the outputs from the simulation platform. It captures the results from simulation experiments in a form that can be directly compared to domain (real world) experimental results, and, potentially, to the domain itself, thereby testing whether the hypothesised low-level mechanisms can result in the observed high-level properties.

The TORCphysics experiments and results are discussed in section 4 section, with different versions of different hypothesised mechanisms matching the domain observations to different degrees.

## 3 Methods – TORCphysics

TORCphysics provides a physical model of supercoiling mediated regulation of gene expression in gene circuits that are sufficiently small that supercoiling propagates effectively instantaneously. Our model accounts for the stochastic transcription initiation for both supercoiling-sensitive and non-sensitive genes (promoters), while also considering the influence of stochastic binding of proteins such as topoisomerases (topoisomerase I and gyrase) and RNAPs. The model quantifies the mechanical impact of these proteins on DNA, specifically addressing supercoiling as changes in the twisting of the double helix; writhe and resulting forms such as plectonemes are not accounted for in the current model. The local structural variations in twist propagate along the DNA, allowing interactions among various bound enzymes and binding sites.

TORCphysics aims to provide the number of RNA transcripts made per second from each gene being considered. It also provides the dynamics of the gene circuit in terms of the interactions between proteins and DNA, transcription events and changes in the superhelical density of the DNA as a function of time. Protein binding to DNA depends on both sequence and local superhelical density, which is parameterised as a pre-processing step. Currently TORCphysics does not consider translation and assumes that there is sufficient ATP for DNA gyrase always to be active and introduce negative supercoils, which corresponds to an environment where the bacteria live in rich media.

### 3.1 Domain Model

#### 3.1.1 Calculating the superhelical density within TORCphysics

DNA topology is described through the linking difference Δ*Lk* = *Lk* −*Lk*_0_, where *Lk* = *Tw* + *Wr* is the linking number defined as the total twist (*Tw*) and writhe (*Wr*) of the topological domain. *Lk*_0_ refers to the relaxed linking number. The superhelical density *σ* can be expressed in terms of the linking difference as:

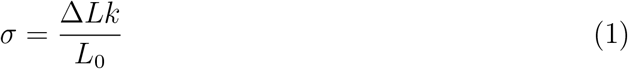

Assuming that changes in topology are exclusively from twist (*Wr* = 0), and that the relaxed linking number corresponds to the total twist of a B-DNA structure of length *L* (in base-pairs), such that *Lk*_0_ = *w*_0_*L*, where *w*_0_ (in rad/bp) is the relaxed twist density of B-DNA, the superhelical density can be reformulated as:

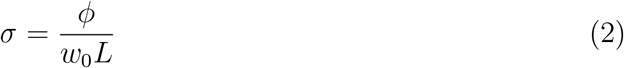

For a topological domain defined by a DNA region constrained at its two ends *x*_*i*_ and *x*_i+1_, the superhelicity within the domain will be isolated from the outside regions. Twisting this region of the DNA by a twist angle of *ϕ_i_* from its relaxed state, the resulting local superhelical density *σ_i_* is given by:

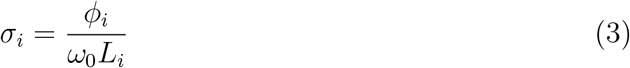

where 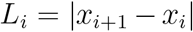 is the length of the topological domain. This assumes that there is no contribution from writhe (i.e., no plectoneme formation) and neglects any non B-DNA motifs such as cruciforms (including stem regions) or single-stranded regions [38].

#### 3.1.2 Topoisomerase activity on DNA

Figure 2a) shows the model we use for describing topoisomerase activity, which is characterised by binding, followed by changes in DNA twist, and finally unbinding from the DNA. We model the binding of topoisomerase I and gyrase on DNA through rate equations 4 and 5. Similar to previous studies [14, 36], we employ sigmoidal functions that modulate the binding rates of topoisomerase I and gyrase as a function of the local superhelical density:

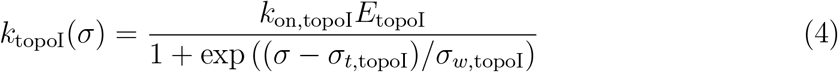

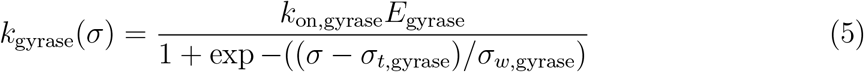

**Figure 2.**
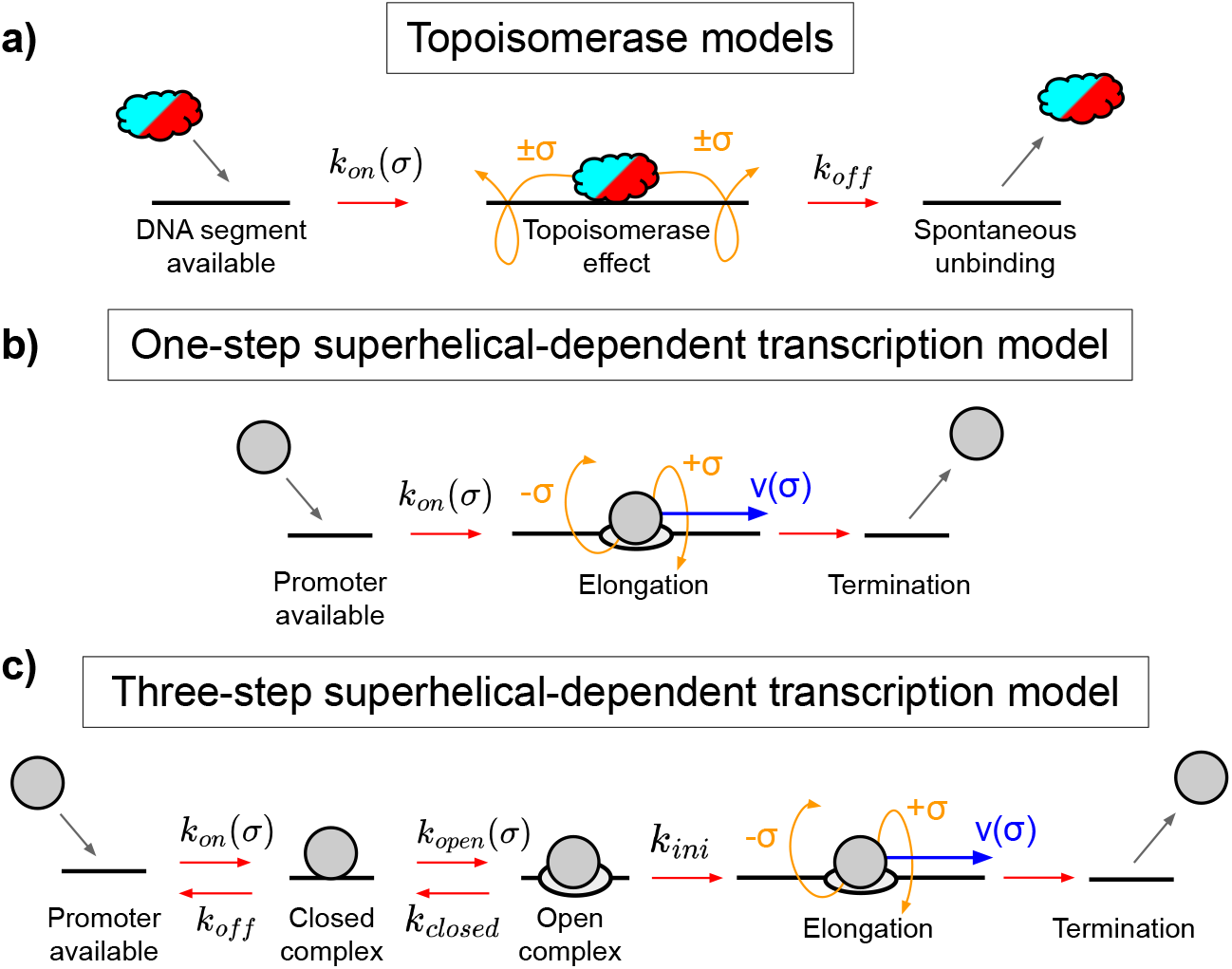
Representation of physical models of enzyme dynamics. a) Models of topoisomerse I (red) and gyrase (cyan) activities in superhelical DNA, where they can bind, modify and unbind the DNA. b) One-step model to describe superhelical-dependant transcription, where elongation starts immediately after the RNAP (grey) binds the DNA. c) Three-step model for describing superhelical-dependant transcrition with several initiation stages before irreversible elongation.

where *k*_topoI_ and *k*_gyrase_ denote the supercoiled-dependent binding rates of topoisomerase I and gyrase respectively (in *s*^−1^), *k*_on_ corresponds to the basal binding rates (in *nM*^−1^*s*^−1^), *E* is the enzyme concentration (in *nM*), and *σ_t_* and *σ_w_* refer to the threshold and width of the sigmoidal function respectively (see table 1). Here we assume the binding of topoisomerase I and gyrase to DNA is determined only by the DNA superhelical density *σ*, and is sequence independent. Depending on the parameterisation of the sigmoidal functions, both topoisomerase I and gyrase can bind to negatively or positively supercoiled regions of DNA.

**Table 1:** TORCPhysics parameters. Binding rates *k*_on_ from Geng et al. [36] were adjusted to match simulation conditions.

| Parameter | Value | Varies | Source |
| --- | --- | --- | --- |
| $\omega_0$ (rad/bp) | $2\pi/10.5$ | No | Intrinsic to B-DNA |
| $\Delta t$ (s) | 1 | No | Simulation |
| $v_0$ (bp/s) | 30 | No | Wang et al. [47] |
| $\gamma$ | $0.157 \pm 0.121$<br>0.01 | No | RNAPTracking section 3.3.3<br>Chatterjee et al. [33] |
| $\kappa$ pN<sup>-1</sup> | 0.5 | No | Sevier et al. [35] |
| $\tau_0$ pN | 12 | No | Sevier et al. [35] |
| $l_{\text{topoI}}$ (bp) | 20 | No | Zhang et al. [42] |
| $l_{\text{gyrase}}$ (bp) | 30 | No | Broeck et al. [43] |
| $l_{\text{RNAP}}$ (bp) | 30 | No | Kang et al. [48] |
| $E_{\text{topoI}}$ (nM) | 17.0 | Yes | Wang et al. [7] |
| $E_{\text{gyrase}}$ (nM) | 44.6 | Yes | Wang et al. [7] |
| $k_{\text{on,topoI}}$ (nM<sup>-1</sup>s<sup>-1</sup>) | $(3.54 \pm 0.042) \times 10^{-4}$<br>$1.06 \times 10^{-4}$ | No | Topoisomerase section 3.3.1<br>Geng et al. [36] |
| $k_{\text{on,gyrase}}$ (nM<sup>-1</sup>s<sup>-1</sup>) | $(2.17 \pm 0.0145) \times 10^{-4}$<br>$0.40 \times 10^{-4}$ | No | Topoisomerase section 3.3.1<br>Geng et al. [36] |
| $k_{\text{off,topoI}}$ (s<sup>-1</sup>) | $0.207 \pm 0.110$<br>0.500 | No | Topoisomerase section 3.3.1<br>Geng et al. [36] |
| $k_{\text{off,gyrase}}$ (s<sup>-1</sup>) | $0.186 \pm 0.076$<br>0.500 | No | Topoisomerase section 3.3.1<br>Geng et al. [36] |
| $k_{\phi,\text{topoI}}$ | $15.11 \pm 3.04$ | No | Topoisomerase section 3.3.1 |
| $k_{\phi,\text{gyrase}}$ | $13.94 \pm 3.42$ | No | Topoisomerase section 3.3.1 |
| $\sigma_{t,\text{topoI}}$ | $0.012 \pm 0.010$<br>-0.040 | No | Topoisomerase section 3.3.1<br>Meyer et al. [14] |
| $\sigma_{t,\text{gyrase}}$ | $-0.059 \pm 0.001$<br>0.0145<br>0.010 | No | Topoisomerase section 3.3.1<br>Geng et al. [36]<br>Meyer et al. [14] |
| $\sigma_{w,\text{topoI}}$ | $(4.25 \pm 1.64) \times 10^{-3}$<br>$12.0 \times 10^{-3}$ | No | Topoisomerase section 3.3.1<br>Meyer et al. [14] |
| $\sigma_{w,\text{gyrase}}$ | $(12.3 \pm 0.2) \times 10^{-3}$<br>$12.2 \times 10^{-3}$<br>$25.0 \times 10^{-3}$ | No | Topoisomerase section 3.3.1<br>Geng et al. [36]<br>Meyer et al. [14] |
| $\sigma_{0,\text{gyrase}}$ | $-0.146 \pm 0.0034$ | No | Topoisomerase section 3.3.1 |
| $\alpha_{E,\text{topoI}}$ | $18.61 \pm 6.78$ | No | RNAPTracking section 4.4 |
| $d_{\text{topoI}}$ (bp) | $448 \pm 40$ | No | RNAPTracking section 4.4 |
| $k_{\text{on, promoter}}$ (s<sup>-1</sup>) | [0.1, 0.33] | yes | Gene architecture section 3.3.2 |
| $k_{\text{off, promoter}}$ (s<sup>-1</sup>) | [0.2, 0.4] | yes | Gene architecture section 3.3.2 |
| $k_{\text{open, promoter}}$ (s<sup>-1</sup>) | [0.01, 0.42] | yes | Gene architecture section 3.3.2 |
| $k_{\text{closed, promoter}}$ (s<sup>-1</sup>) | [0.08, 0.45] | yes | Gene architecture section 3.3.2 |
| $k_{\text{ini, promoter}}$ (s<sup>-1</sup>) | [0.05, 0.42] | yes | Gene architecture section 3.3.2 |
| $\sigma_{e,\text{on, promoter}}$ | [-0.060, -0.045] | yes | Gene architecture section 3.3.2 |
| $\epsilon_{e, \text{promoter}}$ | [0.005, 0.014] | yes | Gene architecture section 3.3.2 |
| $\sigma_{m,\text{on, promoter}}$ | [-0.067, -0.075]<br>-0.042 | yes | Gene architecture section 3.3.2<br>Meyer et al. [14] |
| $\epsilon_{m, \text{promoter}}$ | [0.0089, 0.0093]<br>0.005 | yes | Gene architecture section 3.3.2<br>Meyer et al. [14] |

Once bound, both enzymes can alter the local twist and therefore the superhelical density in a topological domain. To this end, we define the models for the change in twist which are linear with respect to the superhelical density:

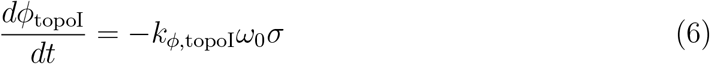

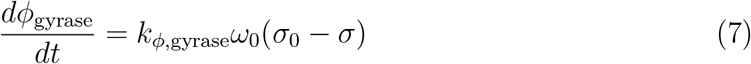

where *dϕ*_topoI_ and *dϕ*_gyrase_ denote the twist angle induced by topo I and gyrase activity respectively, *k_ϕ_* is the rate constant that defines the number of base-pairs by which the DNA is either under or over-twisted per second (bp/s), *ω*_0_ is the twist density (rad/bp) as in equation 3, and *σ*_0_ is the superhelical density threshold at which gyrase can have an effect on the DNA (see table 1). Past this threshold, gyrase is not able to introduce additional negative supercoils into the DNA, and it holds the superhelical density at *σ*_0_ while it remains bound. The angular twist rate can also be expressed as 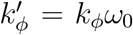 (rad/s); however, we use *k_ϕ_* as this offers an intuitive interpretation in terms of the number of base pairs completely over/under-twisted per second. Specifically, *k_ϕ_* = 1 (bp/s) means that one base pair is fully twisted every second, which corresponds to approximately 0.6 radians per second.

Lastly, we consider the unbinding of topoisomerases to be a spontaneous process, with each enzyme having an associated unbinding rate *k*_off_. Note that topoisomerases can remain bound to DNA for multiple cycles, performing several rounds of enzymatic activity before dissociating.

#### 3.1.3 One-step superhelical-dependent transcription model

Similar to previous studies [14, 27, 36], the one-step model treats superhelical-sensitive transcription initiation as an instant binding/initiation, where RNAPs are recruited to available promoters and transcription immediately starts (see figure 2b). We model this behaviour by employing a sigmoid function, that represents the energy required to melt the promoter, represented by a free energy function *U*_melt_(*σ*):

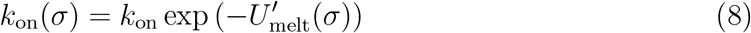

With

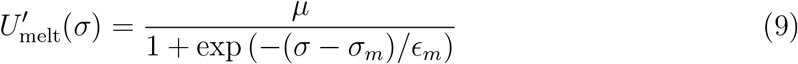

where *k*_on_ denotes the basal rate of RNAP binding, *σ_m_* denotes the threshold for melting, and *ϵ_m_* is the sigmoid width. In practice, rather than directly using the free energy function *U*_melt_(*σ*) (in kcal/mol), we use a dimensionless rescaled version, 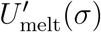, that captures the promoter response to melting via parameters *σ_m_* and *ϵ_m_*. The dimensionless parameter *µ ≈* 2.3 is chosen so that in the inhibitory regime (*σ ≫ σ_m_*) the rate *k*_on_(*σ*) is reduced to 10% of its value, i.e., *k*_on_(*σ → ∞*) = *k*_on_ exp(−*µ*) = 0.1 *k*_on_ (see Supplemental Material Section 3). The free energy *U*_melt_ (and its scaled version 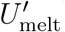) is sequence-dependent and can be parametrised using the SIST algorithm [13] to calculate strand-separation profiles at various superhelical densities. Because of this, binding rate modulation is highly sensitive to GC content, with GC-rich sequences melting more slowly than AT-rich regions, which are more prone to strand separation. The procedure for parameterising *U*_melt_(*σ*) with SIST is detailed in Supplemental Material, Section 3.

Once RNAPs bind their promoters, they automatically transition to the elongation stage, where the RNAP advances along the gene, inducing negative supercoils upstream and positive downstream, until RNAP reaches the termination site. Within this model (equation 10) the build up of supercoils may stall the enzyme jamming transcription. Once the supercoils are alleviated, the RNAP can proceed until it reaches the termination site. These dynamics are captured by the following equations:

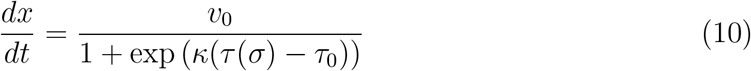

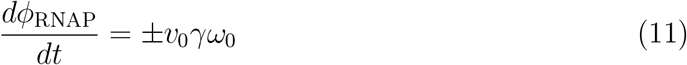

The movement speed of RNAPs is based on the mechanical framework proposed in [35]. The parameter *τ*_0_ = 12 pN corresponds to the stalling torque, and the scaling factor *κ* = 0.5 pN^−1^. The torque *τ* (*σ*) (see figure 2) is modeled using Marko’s elastic model of supercoiled DNA [39] (see Supplemental Material, Section 4 for details). We use an RNAP transcription velocity of *v*_0_ = 30 bp/s, and if the torque exceeds the stalling threshold *τ*_0_, the RNAP stalls. While RNAP velocity is likely variable under realistic biological conditions, we assume a constant velocity for the purposes of this work. The parameter *γ* is dimensionless and quantifies the twist injected by the RNAP per base pair transcribed. The value of *γ* ranges from 0 to 1, where *γ* = 0 corresponds to no twist being imparted on the DNA by the RNAP, implying that the RNAP smoothly rotates around the DNA during elongation, and *γ* = 1 represents a scenario where each base pair transcribed results in complete under- or over-twisting of the DNA. The value of *γ* depends on several factors, such as the viscosity of the surrounding medium, and the size of the transcription complex, which may even include the translational machinery. Here we assume that *γ* is a constant; however, this parameter may vary with transcript length.

Finally, RNAPs advance until they reach a terminator site, where they instantly unbind the DNA.

#### 3.1.4 Three-step superhelical-dependent transcription model

To better capture transcription and its impact on DNA, we propose a three-step model where RNAPs transition through a series of reversible stages of transcription until the elongation phase, as shown in Figure 2c. Upon binding to a promoter, the RNAP forms a closed-complex, the probability of which is modulated by the local superhelical state through an elastic function *G*_elastic_(*σ*), related to the optimal orientation for the promoter [10]. This is followed by a transition to an open-complex, which is governed by the energy required to melt the promoter, represented by the rescaled free energy function 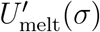. These two key processes are described by:

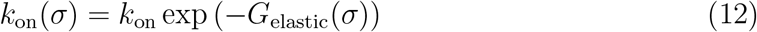

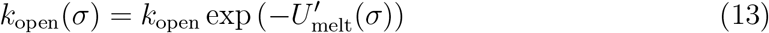

Where

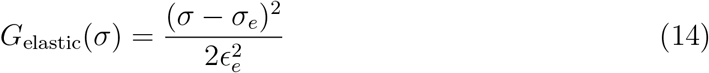

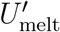 corresponds to equation 9, *k*_on_ and *k*_open_ denote the basal rates of RNAP binding and promoter melting respectively, *σ_e_* and *σ_m_* denote the thresholds for binding and melting, *ϵ_e_* and *ϵ_m_* are the widths of the corresponding distributions. The open complex formation rate *k*_open_ is equivalent to equation 8 in the one-step superhelical-dependent transcription model (see subsection 3.1.3). The elastic function *G*_elastic_(*σ*), which modulates the binding rate *k*_on_(*σ*), is equivalent to the spacer length model proposed by Forquet et al. [10]. While that work employed a specific parameterization of *σ_e_* and *ϵ_e_*, here we use general model for the spacer modulation that can be adapted to an arbitrary promoter sequences and spacer length.

From the open-complex formation, the RNAP then transitions to the irreversible elongation stage according rate *k*_ini_, where the RNAP advances along the gene inducing supercoils. From there on, the model implements the same set of equations for the elongation phase as in the one-step superhelical-dependant transcription model (see subsection 3.1.3 and equations 10-11). Ranges for the promoter-dependent rates and gaussian parameters (e.g. widths *ϵ* and thresholds *σ*) from our calibrations are shown in table 1.

Finally, RNAPs unbind instantaneously upon reaching a terminator site.

#### 3.1.5 Modelling the tracking of RNAP by topoisomerase I

To mimic the behaviour observed in the ChIP-Seq data reported by Sutormin et al. [24], where topoisomerase I tracks the position of transcribing RNA polymerases, we extend the model above (see section 3.1.2) so that the presence of transcribing RNAPs increases the binding rate of DNA topoisomerase I. The expanded binding model for topoisomerase I is of the form:

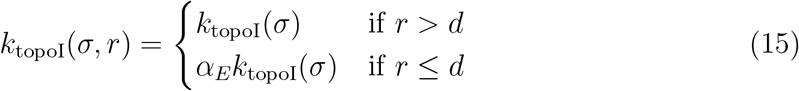

where *r* is the downstream distance between the topoisomerase I binding site and transcribing RNAP (e.g. topoisomerase I follows behind RNAP) *k*_topoI_(*σ*) is binding rate as in equation 4, *d* is an effective distance, and *α_E_* is a multiplier that increases the binding rate of topoisomerase I in the vincinty of RNAP. The binding of topoisomerase I is now influenced by both the superhelical density within the DNA and by the presence of transcribing RNAPs.

### 3.2 Platform model

We now turn to how the domain model above is implemented as a computer simulation, by defining a computational platform model.

#### 3.2.1 The biomacromolecules within TORCphysics

Figure 3a-b illustrates the various biomacromolecules considered within the TORCphysics framework, and the physical abstraction we use to represent each one. The DNA is modeled in one dimension, and since it cannot writhe the only contribution to supercoiling is due to twist. Within this model, the DNA can interact with RNAPs, DNA topoisomerase I, DNA gyrase and NAPs, in a supercoiling dependent manner. DNA-bound molecules are characterised by a position *x_i_* and are associated with a superhelical density *σ_i_*. Protein binding sites can be sequence-specific with well-defined positions (e.g., promoters for RNAPs), or non-specific, allowing proteins to bind anywhere along the DNA (as in the case of topoisomerases). Transcription-supercoiling coupling is captured by considering the supercoiling induced by transcribing RNAPs in accordance with the twin-supercoiling domain model [15], and we assume that all bound proteins act as topological barriers preventing supercoiling diffusion. Although there are no direct interactions between bound molecules modeled within TORCphysics (unless explicitly defined through a specific submodel), they interact indirectly because they all both affect, and are affected by, the superhelical density *σ*.

**Figure 3.**
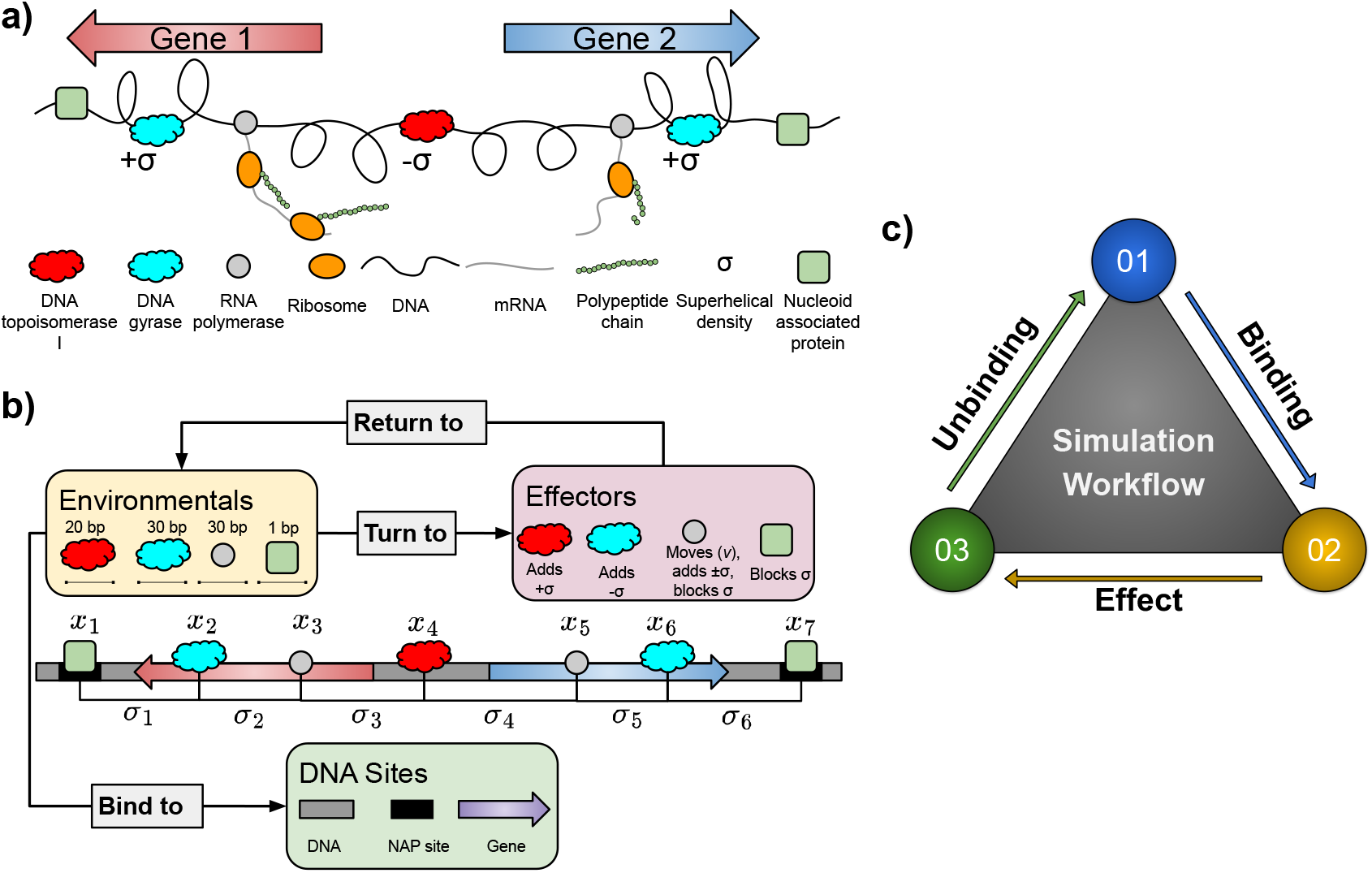
a) Schematic representation of transcription-supercoiling coupling in bacteria using a two-gene system as an example, where RNAPs induce positive supercoils ahead and negative supercoils behind during transcription. b) TORCphysics representation of the transcription-supercoiling coupling in the same two-gene system, where bound proteins are associated with positions *x_i_* and define topological domains characterised by superhelical densities *σ_i_*. c) TORCphysics simulation workflow, where system integration is achieved through cycles of protein binding, effect, and unbinding.

TORCphysics predicts the transcriptional output of a gene as a function of time by considering the evolution of the system in blocks of time Δ*t* (Δ*t* = 1.0 seconds) during which proteins can bind, affect the DNA or unbind. Transcribing RNAPs move fowards with velocity *v* inducing supercoils and acting as moving barriers, while topoisomerases and NAPs are stationary but can modify the supercoiling and block supercoil diffusion. In this work, we do not explicitly model the binding and unbinding of NAPs, but we assume they are always present at the ends in the gene architecture experiments. Since transcription occurs at timescales in the order of minutes/seconds, and supercoiling in the form of twist propagates at the order of *≈* 5kbp/s [40], we assume that supercoiling propagates and equilibrates instantly (within one timestep) in a given region.

Binding and unbinding probabilities of biomacromolecules to a specific DNA site over time Δ*t* are assumed to be independent random events, and so are modelled as Poisson processes [41]:

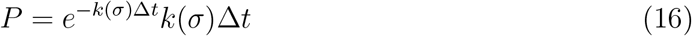

where the rate *k*(*σ*) may be modulated by the local superhelical density *σ*.

#### 3.2.2 Simulation workflow in TORCphysics

In order to quantify the interactions between DNA and biomacromolecules such as RNAPs, topoisomerases and NAPs, our simulation workflow as defined within the TORCphysics framework consists of three stages: *binding, effect* and *unbinding* (see figure 3c), described below. These stages compute the binding and unbinding of molecules in the environment (the *environmentals*) to DNA sites by way of their assigned binding and unbinding models within TORCphysics. When bound, these molecules become *effectors*. Each *effector* has an assigned *effect* model in TORCphysics, which defines how it interacts with DNA. By iterating over these stages, TORCphysics simulates the interaction between the biomacromolecules present and the DNA.

##### *Binding* within TORCphysics

During the *binding* stage of the TORCphysics workflow, we consider the interaction between the DNA and its input environment, which is composed of the *environmentals* (e.g. topoisomerases, RNAPs, NAPs). These *environmentals* all have distinct properties: 1) concentrations *E*, 2) sizes *l*, 3) DNA binding/unbinding behaviour, and 4) specific effector models. The concentrations and sizes are shown in table 1. The current version of TORCphysics does not account for environmental interactions that may influence transcription and protein binding, such as chemical reactions or molecular crowding. All binding models considered in this work are supercoiling dependent. Topoisomerases bind DNA non-specifically according to the TORCphysics model, and so the workflow considers the DNA as split into discrete regions of 20 base pairs for topoisomerase I [42], and 30 base pairs for gyrase [43], and assesses the probability of binding to each using equations 4 and 5, respectively. These sizes were chosen based on crystal [42] and cryo-EM [43] structures of *E. coli* topoisomerase I and gyrase bound to DNA. RNAPs bind only to promoters before they progress along the gene as described in sections 3.1.3 and 3.1.4. Promoters/genes are defined as specific DNA sites having start and end positions, directionalities and particular binding rates/models, such as the one-step or three-step superhelical-dependent transcription models (see equations 8 and 12). NAPs can be modelled to bind to specific DNA sites but have no directionality. The probability of these events are quantified through equation 16.

##### *Effect* within TORCphysics

Once *environmentals* (e.g., topoisomerases, RNAPs and NAPs) have bound, they become *effectors* which have a mechanical impact on the local structure of the DNA: 1) position *x* on the DNA, 2) excluded volume on the DNA due to their size *l* and 3) they define a topological domain with associated twist *ϕ* and supercoiling *σ* (see figure 3b). Within TORCphysics, the mechanical impact of effector *i* is quantified according their change in position along the DNA Δ*x_i_* and change in local twist Δ*ϕ_i_*. Topoisomerases acting as *effectors* do not move along the DNA (Δ*x_i_* = 0), but do change the twist according equations 6 and 7. RNAPs act as topological barriers moving with velocity *v* while injecting negative supercoils upstream and positive downstream (see equation 11). NAPs do not move and do not introduce twist, but they do block supercoil diffusion. Other NAPs, such as LacI, can form DNA-NAP-DNA bridges that create two topological domains. While TORCphysics is capable of modeling this behaviour, it is not explored in the present study.

##### *Unbinding* within TORCphysics

The unbinding of *effectors* is determined by the *unbinding* models. Topoisomerases unbind spontaneously according to an unbinding rate *k*_off_ (see table 1). RNAPs unbind when they reach a transcription termination site, or may unbind before transcription initiation within the three-step transcription model (see section 3.1.4 and figure 2c).

### 3.3 Simulation Platform

Here, we explain how TORCphysics is used to perform computational experiments. We first define the system to be simulated, we then find the biophysical parameters through a calibration process, and we then use the optimal parameter set to perform the final simulation, which provides quantities such as the production rate of RNA from the genetic circuit, the population of biomacromolecules along the DNA and the superhelical densities within the genetic circuits. The calibration processes use a random search algorithm to estimate the best set of biophysical parameters to describe the system. The stochastic nature of the Poisson processes used to represent binding and unbinding events means that it is necessary to perform multiple (typically 100) independent simulations for each set of parameters during the search. Simulations of a typical genetic circuit containing 5 kbp over a simulation time of 2 hr require several minutes to run in real time on a single CPU. Therefore, each computational experiment requires approximately 2,400 CPU hr.

#### 3.3.1 Calibrating the stochastic activity of topoisomerases from experimental data

Here, we calibrate the biophysical models of the stochastic activity of topoisomerases on superhelical DNA (see section 3.1.2). Wang et al. [7] used a fluorescent marker to quantify how fast topoisomerase I and gyrase relax and negatively supercoil DNA respectively, and show that both topoisomerases exhibit classic Michaelis-Menten kinetics. In Supplementary Material Section 5, we show how we construct reference curves for the change in DNA superhelical density *σ*_kinetic_ with time for the two topoisomerases, both individually and in combination (shown in figure 4). To discretise these kinetic processes in time for use in TORCphysics models, we simulated an equivalent plasmid (2757 bp), topoisomerase concentrations (see table 1) and timescale *T* (*T* = 500 seconds) as used by Wang et al. [7] to obtain a DNA superhelical density *σ*_TP_. We tested 8000 random parameter sets for equations 4–7, and for each performed an ensemble of 100 repeat simulations, calculating an average superhelical density *σ*_TP_ per ensemble. We then selected the set of parameters that best fit the reference curves *σ*_kinetic_ by minimising the objective function:

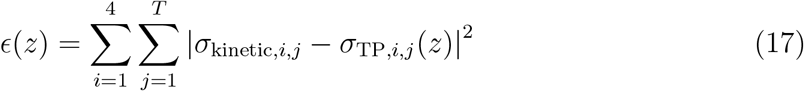

**Figure 4.**
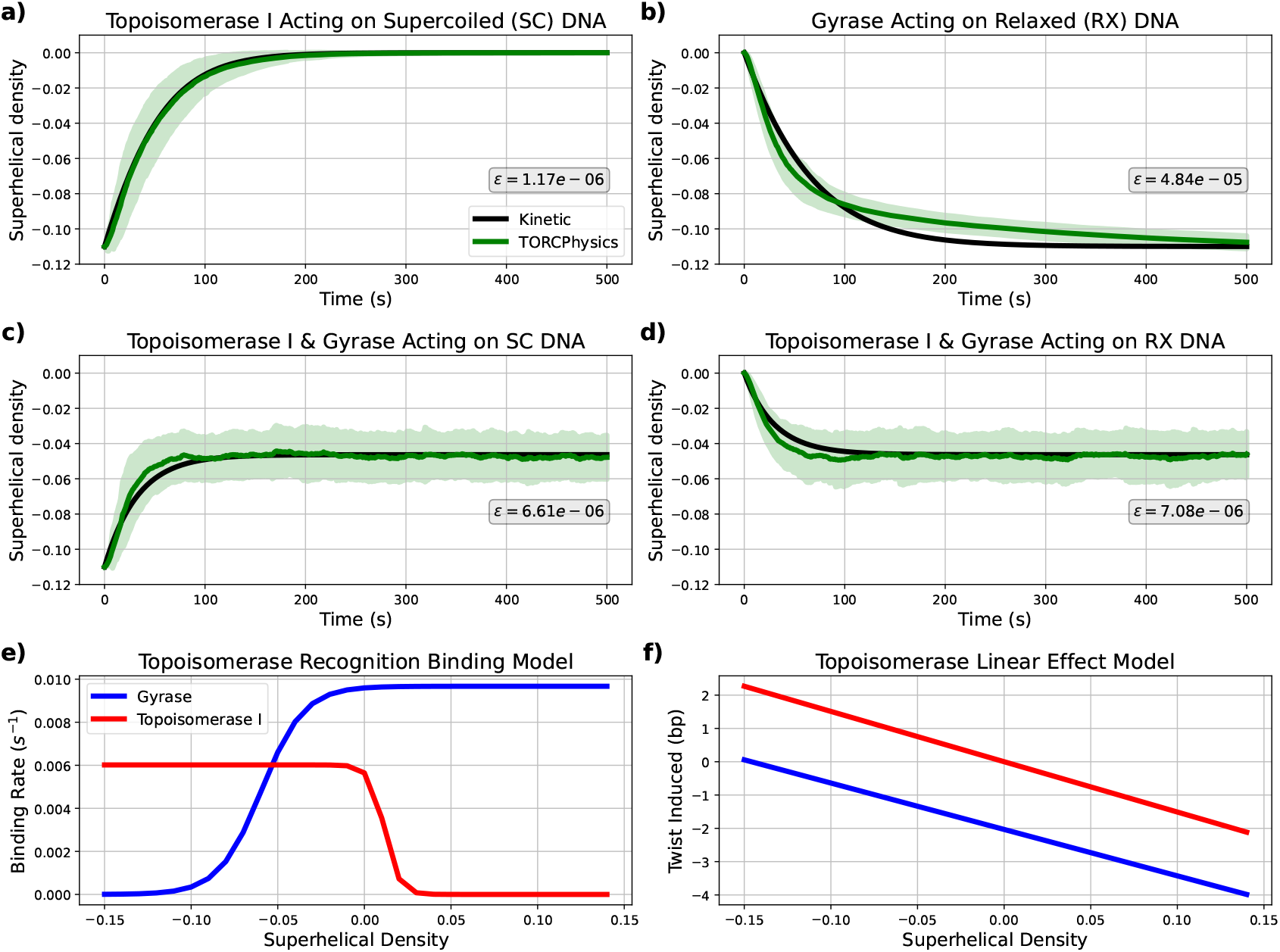
Stochastic topoisomerase activity parameterised from experimental data [7]. TORCPhysics results using the best parameter set from the calibration process described in section 3.3.1 (green curves) compared with kinetic curves inferred from experimental data (black curves) across different scenarios: a) Topoisomerase I acting on negatively supercoiled DNA, b) Gyrase acting on relaxed DNA, c) Both topoisomerases acting on negatively supercoiled DNA, and d) Both enzymes acting on relaxed DNA. Shaded green areas represent the standard deviation obtained from multiple simulations. Superhelicaldependant binding rates (e) from equations 4 and 5, and rates of twist (f) from equations 6 and 7.

where *z* denotes to the parameter space, *j* labels the time data points (1 to *T*), *i* labels four scenarios: (1) topo I acting on negatively supercoiled DNA, (2) gyrase on relaxed DNA, (3) both enzymes acting on supercoiled DNA, and (4) both enzymes acting on relaxed DNA (see figure 4). The parameter space *z* includes 11 parameters: binding rates *k*_on_, sigmoid parameters *σ_w_* and *σ_t_*, unbinding rates *k*_off_, twist rates *k_ϕ_* and the maximal superhelical density that gyrase can impose *σ*_0_. Supplemental Figure S1 shows the distribution of losses, confirming that the calibration process successfully converged to acceptable solutions for the purposes of this work.

#### 3.3.2 Calibrating promoter supercoiling responses from genetic architecture experiments

We now calibrate promoter activity using the transcription models presented in sections 3.1.3 and 3.1.4, by matching TORCPhysics outputs with the changes in gene expression rates as a function of gene architecture measured by Junier et al. [18]. Those experiments used a plasmid containing two genes coding for fluorescent proteins, where one is flanked by two LacI binding sites. When bound, the LacI-looping protein isolates the inner gene. That experiment studied how the ratio of fluorescence changes when different DNA fragment lengths are inserted between the upstream barrier and the promoter of the enclosed gene. The experiment investigated three promoters with different strengths, ranging from weak to strong (see promoter sequences in Supplementary Table S3). For all three promoters, they observed that gene expression increases as the upstream barrier is positioned further away.

To evaluate the ability of TORCphysics to reproduce those experimental observations, we performed simulations for circuits composed of flanking barriers (modelled as NAPs) with a single gene in between, where the shortest distance between the upstream barrier and the promoter is 101 bp and the longest 5051 bp, and the gene length is 900 bp. The stochastic activity of topoisomerases was modelled using the parameterisation in section 3.3.1. To mimic transcription we consider three physical models of increasing complexity:

- V0: One-step transcription model (section 3.1.3).
- V1: Three-step transcription model (section3.1.4).
- V2: Three-step transcription model with topoisomerase I RNAP tracking (sections 3.1.4 & 3.1.5).

Since both the one- and three-step transcription models have expressions that depend on the free energy required to melt DNA (*U*_melt_), which depends on both the DNA sequence and the superhelical levels, we first computed the stress-induced DNA duplex destabilization profiles (SIDD) [12] for the three promoter sequences before beginning the calibration process. To achieve this, we added a flanking sequence of 250 G/C base pairs on each side to each promoter sequence. Using the SIST algorithm [13], we calculated the free energy required to melt each promoter at various superhelical densities (see Supplemental Figure S2). Finally, we fitted a sigmoidal function to model open complex formation as a function of the melting energy (see Supplemental Figure S3).

We then perform a calibration process for the three promoters, running 60 simulations lasting 1.5 hours of simulation time, for each upstream distance *i*. From these, we calculate the susceptibility *s*, defined as the expression rate *k* divided by the expression rate at a reference upstream distance *s_i_* = *k*_*i*_/k_*i*=ref_. We select the reference upstream distance 250 bp for comparison with Junier’s data [18]. We conducted 300 random tests for V0 and 1300 for V1 and V2, evaluating each test performance according to the objective function:

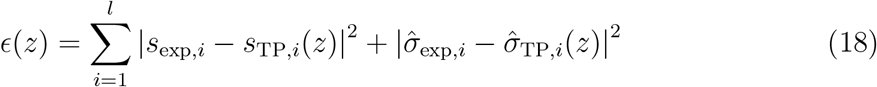

where *s*_exp*,i*_ denotes the experimental averaged susceptibility for the upstream distance *i, s*_TP*,i*_(*z*) is the corresponding simulated susceptibility, 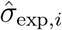 is the experimental standard error and 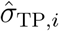 is the simulation standard error. The parameter space *z* includes the promoter rate *k*_on_ in case of V0 (one parameter to fit), while for V1 and V2 it includes the promoter rates of the three-step transcription model, as well as the width *σ_e_* and threshold *ϵ_e_* from the elastic function *G*_elastic_ (seven parameters to fit, see Equations 12, 13 and Figure 2c). Finally, for V0 and V1 we independently calibrate *γ*, but for V2 the twist injection rate is obtained through the topoisomerase I tracking protocol described in section 3.3.3.

For V0, the optimal solutions are found with *γ* = 0.05, for V1 they are found with *γ* = 0.1. Supplemental Figure S4 presents the distribution of losses across the three physical models (V0, V1, and V2), demonstrating that the calibration processes successfully converged to acceptable solutions for the purposes of this work.

#### 3.3.3 Calibrating the tracking of RNAP by Topoisomerase I from ChIP sequencing data

The ChIP-Seq data obtained by Sutormin et al. [24] demonstrates that DNA topoisomerase I tracks transcribing RNAPs in *E. coli*. To assess the importance of this phenomenon on gene regulation, we have included this interaction within TORCPhysics using the three-step transcription and the tracking topoisomerase I models from sections 3.1.4 and 3.1.5, respectively.

To mimic the ChIP-Seq analysis, we simulated a 6000 bp sequence (this is the typical size of a topological domain found in *E. coli* [44]) containing an averaged sized gene of 1000 bp [45] (see figure 5). We assume the DNA ends are constrained by topological barriers, and that the superhelical density can be modified only by the action of topoisomerases. We used the parameters for topoisomerase activity and the initial superhelical density (-0.046) obtained previously (see section 3.3.1) and ran each simulation for 1000 seconds (16.6 minutes of simulation time).

**Figure 5.**
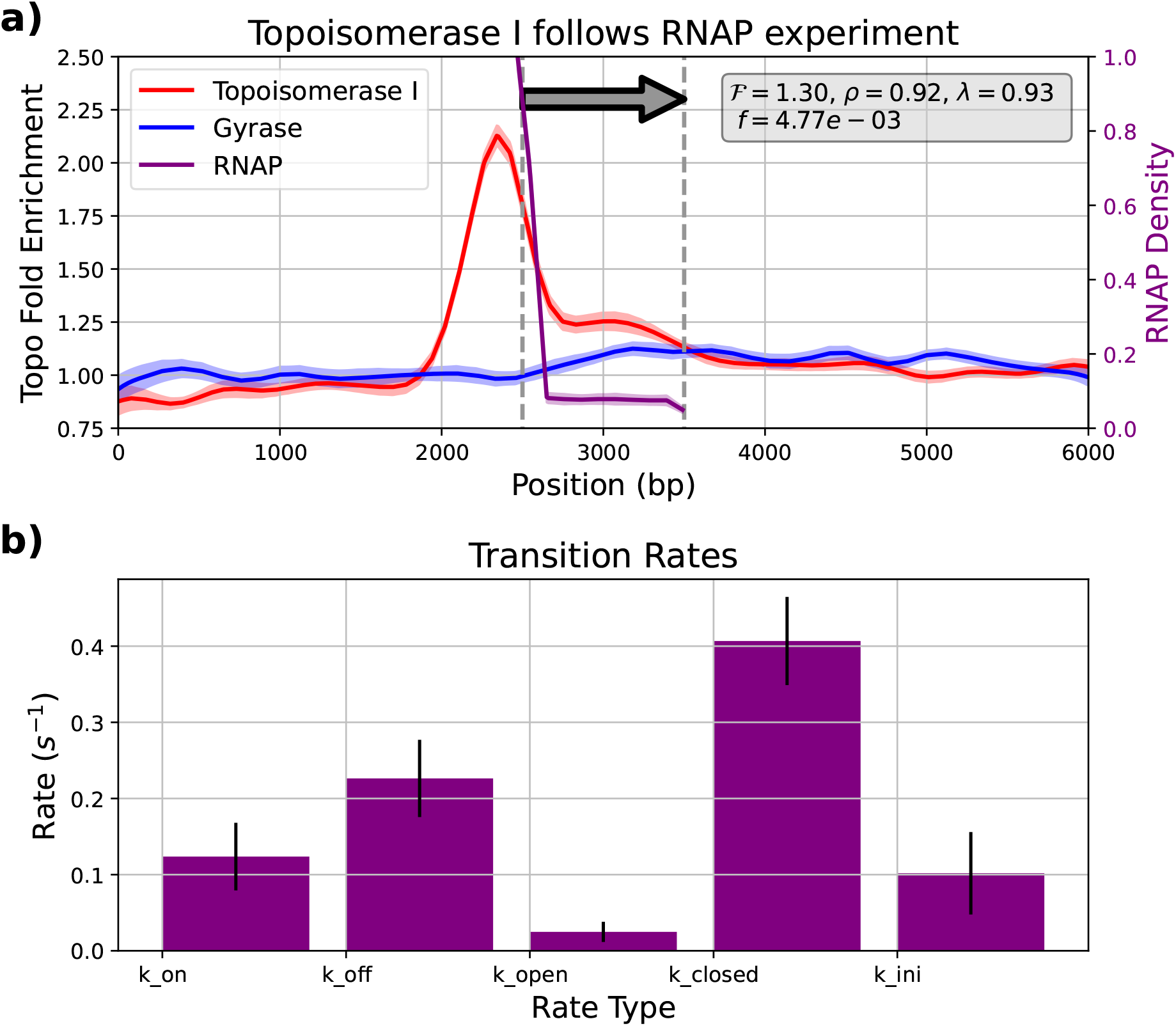
a) Fold enrichment profiles for topoisomerase I (red), gyrase (blue), and RNAP normalised density (purple), obtained from the topoisomerase I tracking RNAP calibration process described in section 3.3.3, using the optimal set of parameters. The grey arrow represents the transcription unit (TU). Parameter ℱ denotes the mean fold enrichment of topoisomerase I within the TU, *ρ* represents the correlation coefficient between RNAP and topoisomerase I positions, *λ*(*z*) indicates the correlation between RNAP density in TORCphysics and ChIP-seq data from Sutormin et al. [24], and *f* represents the loss function (see equation 19). b) Transition rates used in the three-step superhelicaldependent transcription model (see Figure 2). Bars indicate average values, and error bars represent standard deviations across the top 5% of best calibration sets.

We quantify the relationship between the transcribing RNAPs and DNA topoisomerase I by calculating the Pearson correlation coefficient *ρ*. Using ChIP-Seq [24] to deduce the two enzyme position densities within the transcribing unit (TU) gives *ρ* = 0.94 (see Supplemental Figure S5). The fold-enrichment compares ℱ the topoisomerase I activity in the presence of transcribing RNAPs with the case where there is no transcription. Calculating an average over the TU from the ChIP-Seq data gives ℱ = 1.24 (see Supplemental Figure S5). We also calculate the correlation coefficient *λ* between the simulated RNAPs and the experimental RNAP signal within the TU to parametrise the action of transcribing RNAPs using the three-step transcription model (see section 3.1.4). We implement the following four step calibration protocol:

- Step 1: We start with a reference system where transcription is turned off and only topoisomerase I and gyrase interact with DNA by way of the stochastic topoisomerase model 3.1.2. We launch a set of 8 simulations and calculate position densities for both enzymes. Each simulation runs for 1 hour of simulation time.
- Step 2: Transcription is now turned on, and we perform 3000 random tests with different parameterisations to calculate the objective function *f* (*z*) (see equation 19). For each test, we run 28 sets composed of 8 simulations each, which are combined to give 28 histograms describing the position densities for RNAP, topoisomerase I and gyrase. The fold-enrichments ℱ are computed and averaged within the TU, and the mean position densities are obtained per molecule (RNAP, topo I and gyrase).
- Step 3: The mean densities are used to compute the correlation coefficient *ρ* between RNAP and topoisomerase I.
- Step 4: The calibration process is complete when the parametrisation set that minimizes the objective function has been determined.

The objective function minimsed during the calibration is :

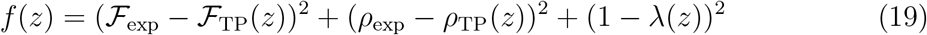

where *z* denotes the parameter space to be calibrated, ℱ _exp_ = 1.24 is the mean foldenrichment of topoisomerase I within the TU measured experimentally, ℱ _TP_(*z*) is the fold-enrichment calculated from simulations, *ρ*_exp_ = 0.94 is the experimental correlation coefficient between RNAPs and topo I, *ρ*_TP_(*z*) is the correlation coefficient from the simulations, and *λ*(*z*) is the correlation between the experimental RNAP density and the density within the TU region obtained from simulations.

The parameter space *z* comprises the effective distance *d* and multiplier *α_E_* from the topoisomerase I tracking model (equation 15), the rate of twist *γ* induced during transcription (equation 11), and the promoter rates *k*_on_(*σ*), *k*_off_(*σ*), *k*_open_(*σ*), *k*_closed_, and *k*_ini_ for the three-step transcription model (see Figure 2c). For the elastic function *G*_elasti_ used in the three-step transcription model, we select the values *σ_e_* = −0.06 and 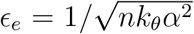 (see Supplemental Section 6). This parametrization is equivalent to the spacer length model proposed by Forquet et al. [10]. For the melting energy function *U*_melt_, we use *σ_m_* = −0.042 and *ϵ_m_* = 0.005, which correspond to those used in the thermodynamic model from El Houdaigui et al. [14] for the *pelE* promoter [46]. We use this fixed parametrization to represent the supercoiling response of an average bacterial promoter. The distribution of losses shown in Supplemental Figure S6 demonstrates that the calibration process presented here converged to acceptable solutions.

## 4 Results and Discussion

### 4.1 Modeling the stochastic activity of DNA topoisomerase I and DNA gyrase on supercoiled DNA

In bacteria, the activity of topoisomerase enzymes is crucial for maintaining an adequate negative superhelical level [2–4]. Here, based on the kinetic parameters measured by Wang et al. [7] in *E. coli*, we have derived a discretised biophysical model for the stochastic activity of topoisomerases on supercoiled DNA.

Figure 4 shows the superhelical densities inferred from the steady-state kinetics [7], compared with the average superhelical densities obtained from simulations using the optimal parameter set from the calibration process (see Table 1). Overall, there is strong agreement between the experimentally determined kinetics and the TORCphysics results, with relatively small mean squared errors on the order of *ϵ ≈* 10^−5^ for all four cases. For topoisomerase I acting on supercoiled DNA, the model more accurately captures the relaxation curve, with a very small error (*ϵ ≈* 10^−6^; see Figure 4a). In contrast, for gyrase acting on relaxed DNA (Figure 4b), the model generally reproduces the kinetic behaviour but tends to overestimate the amount of torsion (negative supercoils) introduced. Topoisomerase I acts faster than gyrase, completely relaxing hyper-negative supercoils within 100 seconds, while gyrase takes approximately 200 seconds to negatively supercoil the DNA. In the combined scenarios where both enzymes act on supercoiled/relaxed DNA (Figure 4c-d), the model performs well over longer timescales, although there is an overestimation of the superhelical in the first 100 seconds before the steady-state plateau is reached.

Figures 4e-f show the binding rates and twists induced by DNA topoisomerase I and gyrase obtained from the calibration process. For the grid sizes, enzyme concentrations and kinetic rates in table 1, and at superhelical densities where they are fully active, we observe that both enzymes bind to the DNA at an approximate rate of one every 2 seconds, and remain bound for around 5 seconds before unbinding. A binding event every 2 seconds aligns with single-molecule experiments of DNA gyrase activity *in vivo* [49] and is 3 to 5 times more frequent than values used in previous stochastic models [36] (see Table 1). In terms of unbinding, our predictions are about five times higher than in previous models calibrated for *E. coli* [36]. However, our values are consistent with single-molecule experiments of gyrase activity around the replisome [49]. The set of estimated parameters indicate that both DNA topoisomerase I and gyrase maintain significant presence on the DNA across physiological superhelical densities *σ* = [− 0.11, 0.05], and that both enzymes exhibit similar behaviour in terms of binding and unbinding rates, as well as catalytic activity *k_ϕ_* (see Table 1). Experimental and theoretical studies have also reported that both enzymes have comparable activity [7,14,36] and propose sigmoidal functions to model activity. Although the threshold parameters *σ_t_* used are different, the width parameter *σ_w_* is similar across these studies. For instance, El Houdaigui et al. [14] report a width twice as broad as in TORCphysics, while Geng et al.’s model [36] yields an almost identical width parameter for gyrase (see Table 1). However, Meyer’s model differs from TORCphysics topoisomerases by acting continuously in every topological domain.

Figure 6 shows the number of bound enzymes throughout the simulation, indicating that each system reaches a steady state. For DNA topoisomerase I, we observe that the DNA remains relaxed in spite of the presence of the enzyme, as expected as topoisomerase I primarily relieves superhelical stress [2] (see Figures 6a and 4a). In contrast, gyrase activity decreases as the DNA becomes hypernegatively supercoiled (see Figures 4b and 6b). The steady-state curves when both enzymes are fully active (Figures 6c-d) show that approximately 2 DNA topoisomerases and 2.5 gyrases are bound to the DNA on average and maintain a steady-state superhelicity of −0.046, which is consistent with *in vivo* observations [50]. This behaviour suggests that at least one enzyme of each type is present per 1kb of DNA, which aligns with the average gene size in prokaryotes [45].

**Figure 6.**
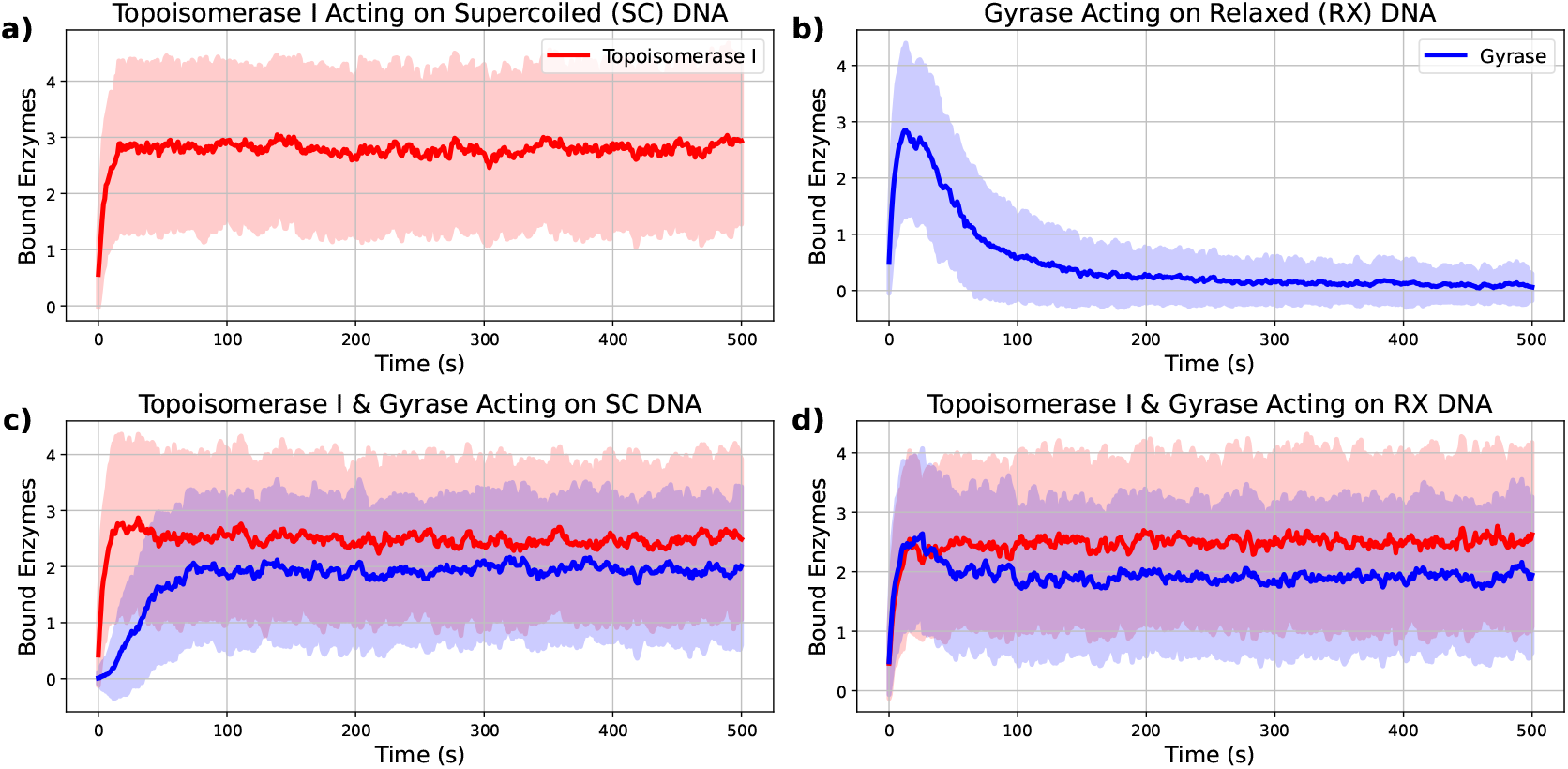
Steady-state curves resulting from the stochastic topoisomerase activity experiment described in section 3.3.1, using the best parameter set. The plots show the average number of topoisomerase I (red) and gyrase (blue) molecules bound to DNA over time, across different scenarios: a) Topoisomerase I acting on negatively supercoiled DNA, b) Gyrase acting on relaxed DNA, c) Both topoisomerases acting on negatively supercoiled DNA, d) Both enzymes acting on relaxed DNA. Shaded areas represent standard deviations obtained from multiple simulations.

Based on these observations, we consider the calibration procedure for obtaining the kinetic parameters for topoisomerase I and gyrase to be sufficiently accurate that we can justify the introduction of additional complexity into the TORCphysics models. In our gyrase model, neither ATP nor ADP are considered, and a more sophisticated treatment that considers ATP/ADP ratio as well as ATP hydrolysis could potentially improve our description of gyrase activity in future work. Here, we assume that ATP is always available to gyrase, allowing it to introduce negative supercoils and relax positive ones. In the absence of ATP, gyrase can relax negatively supercoiled DNA [6]; this behaviour could also be modeled with TORCphysics using a similar calibration process, but with experimental data reflecting those specific conditions.

### 4.2 Impact of gene architecture on expression within the onestep transcription model (V0)

DNA supercoiling and gene expression are intimately linked. Boulas et al. [18] investigated this relationship *in vivo* within the simple genomic context of a system consisting of a single gene enclosed within a topological domain. Their experiments showed that gene expression is strongly influenced by the distance between the promoter and the upstream topological barrier, while their biophysical model proposed that topoisomerase I represses initiation and facilitates elongation. Here, we simulate this system with TORCphysics using the calibrated topoisomerase models and the one-step transcription model.

Figure 7a-c shows the susceptibility calculated as a function of the distance between the promoter and the upstream barrier following calibration (3.3.2). Of the three models investigated, V0 gives the least favourable comparison with the experimental data. Analysing the gene expression rate versus the objective function *f* (*z*) (see figure 7a-c), we see that the expression rate is only one transcript approximately every 7 minutes. However, Figure S7d shows that the optimised RNAP binding rates are around one event per minute.

**Figure 7.**
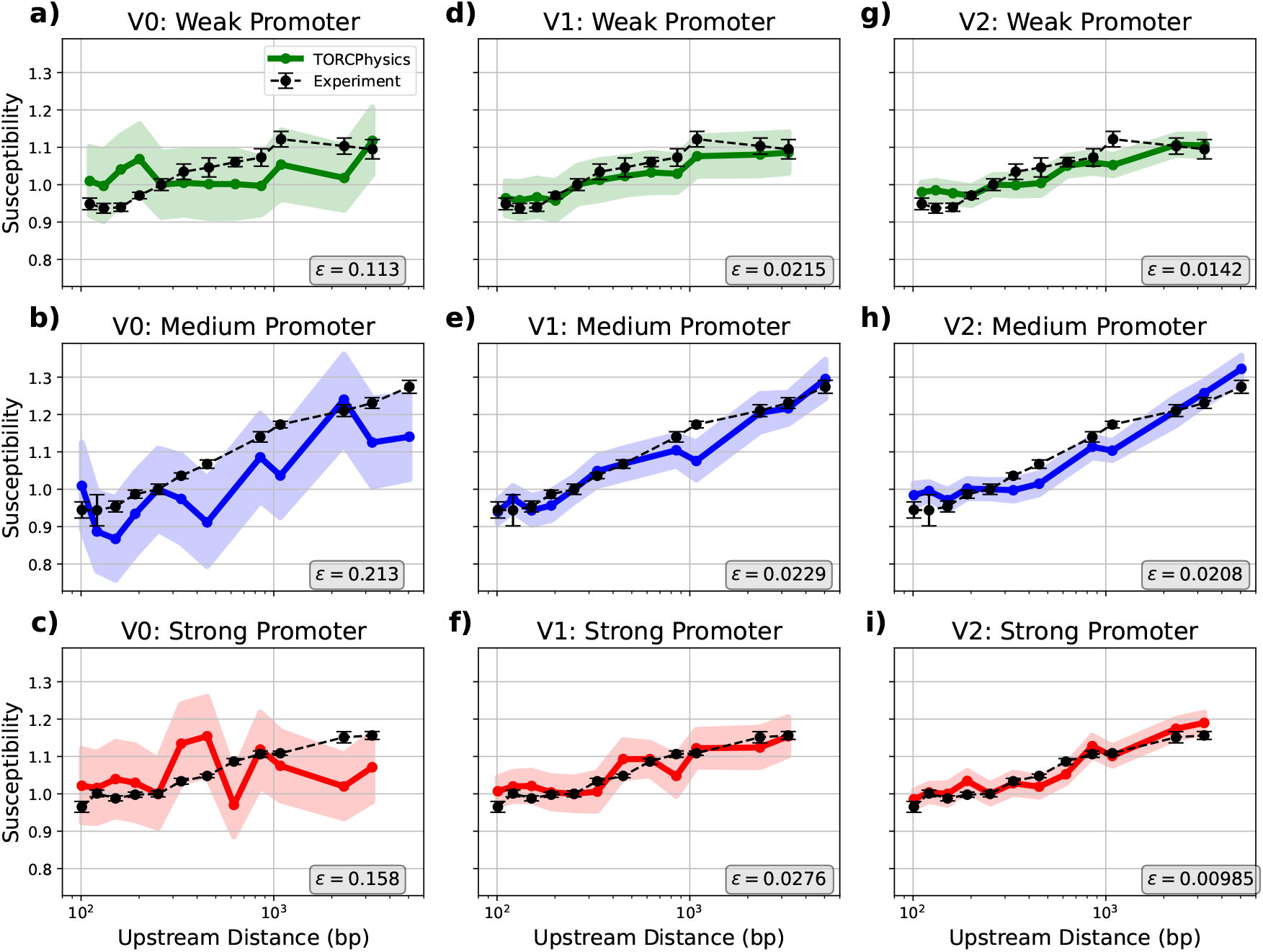
Averaged susceptibility calculated from TORCphysics as a function of upstream barrier distance for the weak (green), medium (blue), and strong (red) promoters, compared with the experimental susceptibility measured by Boulas et al. [18] (black dashed line). The TORCPhysics results were obtained by the calibration process for three models V0 a-c, V1 d-f, and V2 g-i, described in section 3.3.2. Standard errors are represented as shaded areas for the TORCphysics simulations and as error bars for the experimental data.

To understand this discrepancy, we analysed the average number of bound topoisomerases and RNAPs (see figure 8a-c). For both topoisomerase I and gyrase we see a higher number of bound enzymes as the distance between the promoter and the barrier increases as there is more space available for these molecules to bind. However, the number of elongating RNAPs saturates, presumably because the high levels of supercoiling have jammed transcriptional activity. These results indicate that the one-step transcription model is not capable of representing the experimental system.

**Figure 8.**
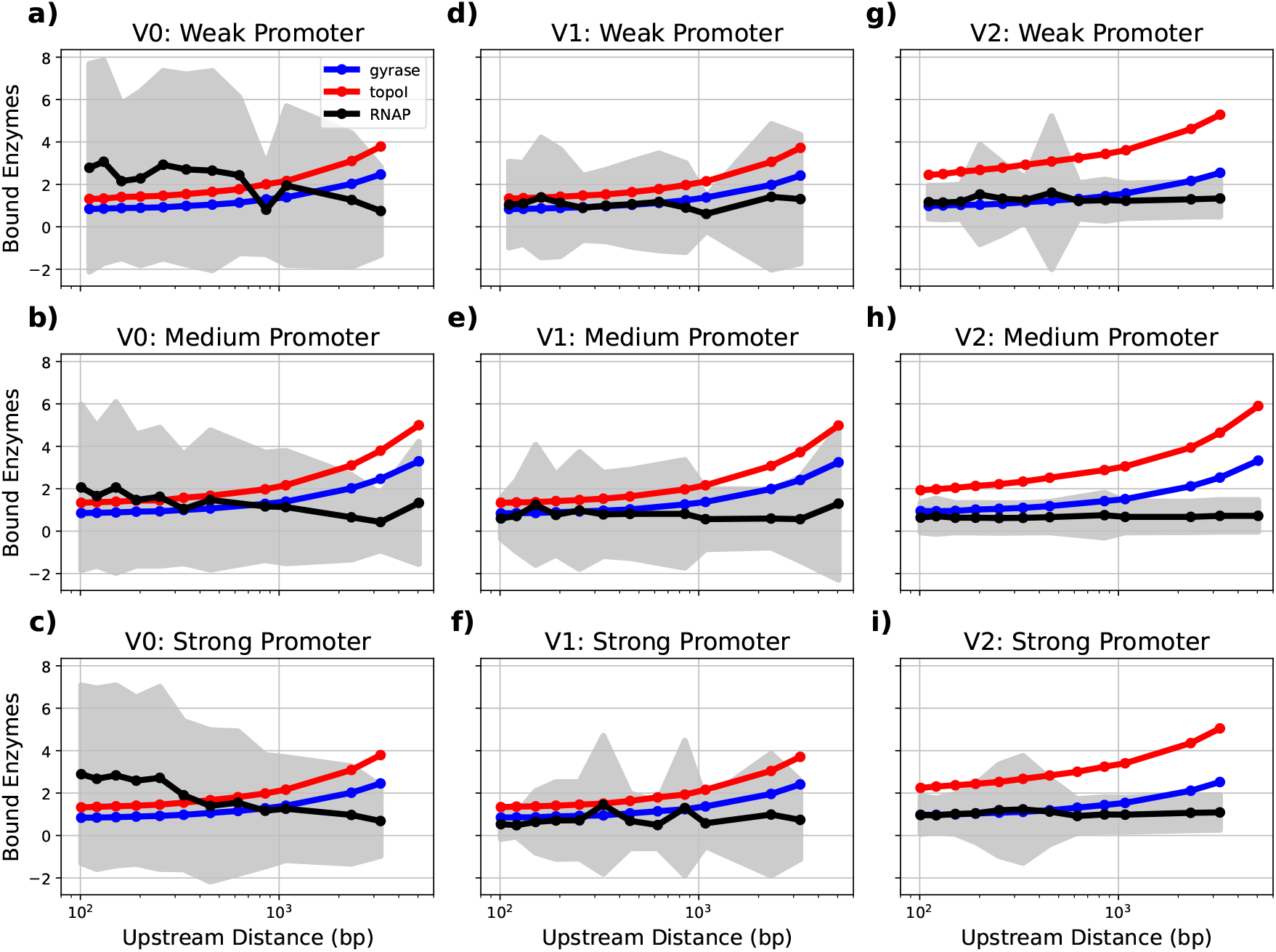
Average number of bound enzymes as a function of the distance between the upstream barrier and the promoter, using the optimal promoter parameterisations for the three models: V0, V1, and V2. Topoisomerase I is shown in red, gyrase in blue, and RNAPs in black, with standard deviations represented as shaded areas.

### 4.3 Impact of gene architecture on expression within the three-step transcription model (V1)

We now consider how the expression rate changes with distance between the promoter and the topological barrier using the three-step transcription model within TORCphysics, as shown in figure 7d-f. As well as improved agreement with the experimental data, we see from figure 9d-f that higher expression rates of 1 event every 3 minutes are obtained. Figure S7e shows the rates obtained for the optimal calibration, and demonstrates that promoter activity depends on a complex balance of kinetic properties, consistent with previous studies [51].

**Figure 9.**
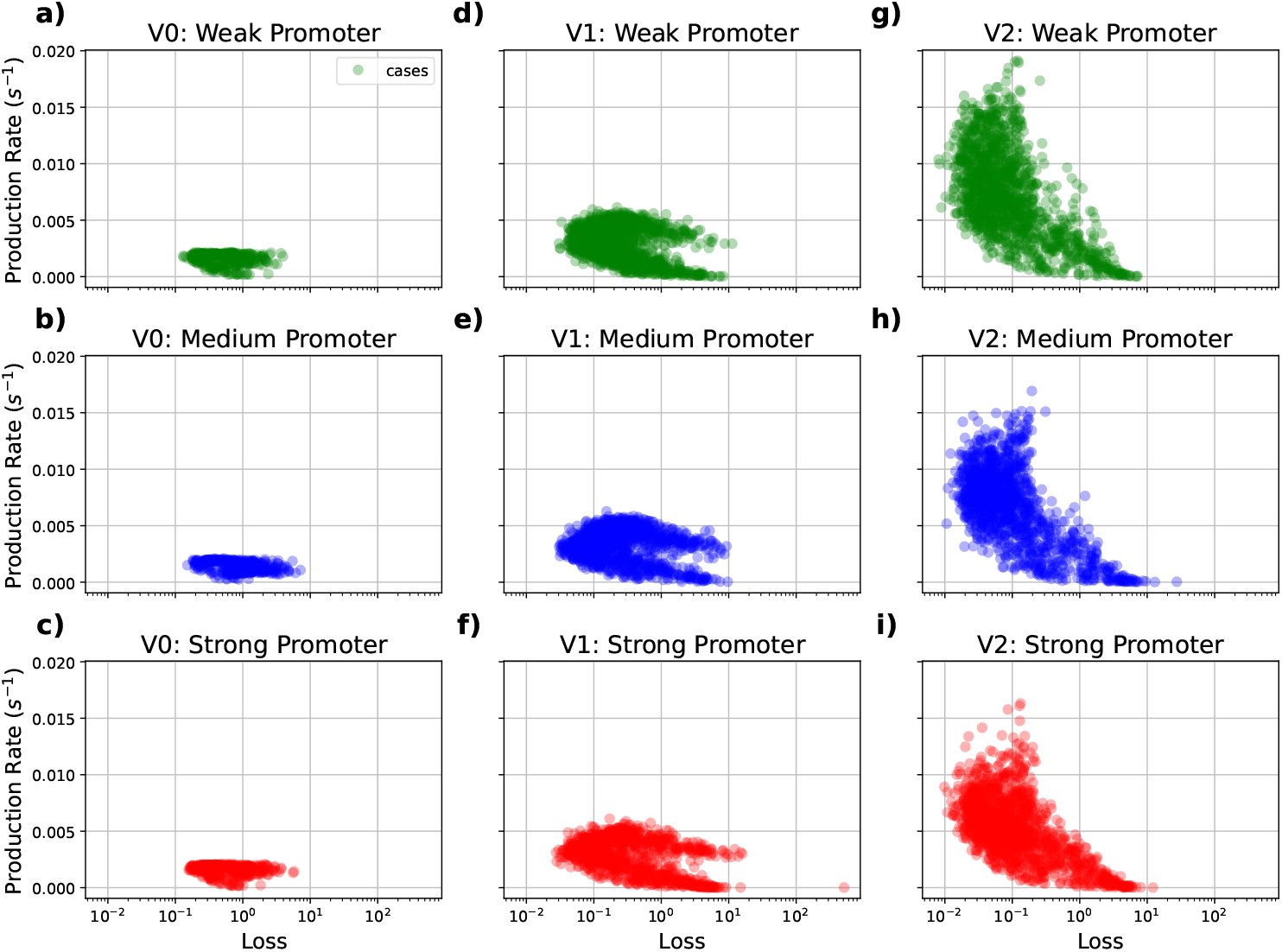
Averaged production rates from all parameterisation sets tested during the calibration process described in section 3.3.2 for the three model sets: V0, V1, and V2. Values for the weak promoter are highlighted in green, medium in blue and strong in red.

While the number of bound topoisomerases is comparable for V0 and V1, the number of transcribing RNAPs for V1 is lower (on average 1 RNAP for V1 compared to 3 for the V0 model), which is surprising given that the transcription rate is higher for the three-step transcription model (see figure 8d-f). While the one-step transcription enables faster initiation events, the number of elongating RNAPs saturates. However, the three-step model allows the RNAP to pause during initiation, which prevents jamming and leads to a higher gene expression rate. Nevertheless, the expression rate is still only around twice as fast for the three-step compared to the one-step transcription model.

### 4.4 RNAP tracking by Topoisomerase I

Recent ChIP-Seq studies have shown that *E. coli* DNA topoisomerase I directly interacts with transcribing RNA polymerase [22–24]. To produce a model of this interaction for use within the TORCPhysics framework, we built an average topological domain including a single central gene, and implemented the protocol in 3.1.5. Figure 5a shows the computed fold enrichment for both topoisomerase I and gyrase, as well as the RNAP densities calculated from the simulations using the optimised parameter set. Topoisomerase I shows an enrichment of around 1.3 along the TU, compared to the ChIP-Seq data value of 1.4 reported in [24]. For the correlation coefficient *ρ* between topoisomerase I and RNAP, the model obtains nearly 92%, compared to the experimental value of 94%. Lastly, the models shows a 93% correlation between the simulated RNAP densities and the RNAP ChIP-Seq data (see Supplemental Figure S5).

The effective distance *d* and multiplier *α_E_* obtained from the calibration procedure in section 3.1.5 indicate that the RNAP position enhances the binding of topoisomerase I by 18 fold, and this effect extends to between 400 and 500 base-pairs behind bound RNAPs (see table 1). However, the maximum Fold-Enrichment of topoisomerase I is only around 2, because the topoisomerase I binding rate is so low (see figure 5a). The estimated rate of twist injection *γ* induced by elongating RNAPs extracted from our calibration procedure ranges from 0.03 to almost 0.30, which are both within the expected theoretical values (see table 1). Previous studies have used lower values of *γ* = 0.01 [33].

Figure 5b shows the promoter rates for the three-step transcription models that resulted from the optimization process in the presence of RNAP tracking by topoisomerase I. The rate-limiting step is open-complex formation *k*_open_ (which is approximately 1 event every 35 seconds). The balance of these rates indicates that there are successive rounds of failed initiation and short-lived intermediate states before successful transcription initiation, consistent with previous observations [52–54]. However, open-complex formation is not always a limiting step, as each promoter has its own unique kinetic properties, which together determine promoter strength and response [51].

### 4.5 Impact of gene architecture on expression in the presence of RNAP tracking by topoisomerase I (V2)

Finally, we use the calibrated three-step transcription model including RNAP tracking by topoisomerase I to calculate the gene expression rates as a function of distance between the promoter and the topological barrier, for comparison with the experimental data from Boulas et al. [18]. Figures 7g-i show the susceptibility as a function of the upstream distance for the three promoters. The average error calcuated across the three promoters is reduced from 0.161 for V0 and 0.024 for V1 to 0.015 for the V2 model.

Figure 9g-i shows that the averaged gene expression rates is increased to 1 transcript per minute, which is a factor of 3 improvement compared to V1. The number of bound topoisomerase I enzymes increases by 1 (figure 8g-i), while the number of bound RNAPs actually decreases in comparison with V0 and V1, and gyrase activity remains unchanged. These results suggest that the RNAP tracking mechanism included in V2 provides one extra bound topoisomerase I, which in turn results in increased RNAP elongation where rounds of transcription are characterized by a single RNAP transcribing one at a time.

Taking into account all these features reveals a complex network of mechanisms. Equation 3 implies that negative supercoils accumulate faster when the transcribing RNAPs are closer to the topological barrier. These negative supercoils can both promote transcription (by melting the promoter) or stall transcribing RNAPs due to the increasing levels of torsion. Including the tracking of RNAP by topoisomerase I within the model faciltates the removal of supercoils when RNAP is stalled. When the upstream distance from the promoter to the topological barrier is larger, the build up of negative supercoiling is slower. RNAPs can advance longer distances before stalling, and topoisomerases have more accessible space for binding and removing the excess supercoiling. This results in a higher susceptibility and greater gene expression rates. When these effects are combined, it is possible for a gene to both promote and repress itself through its own transcription, depending on the size of the topological domain.

Our simulations therefore suggest an alterative potential mechanism for the results reported by Boulas et al. [18], who proposed that transcriptional repression was driven by topoisomerase I activity. Instead, our model indicates that the dynamic interaction between stochastic topoisomerase I activity and gene expression can create a self-regulating system. Although previous studies have shown that topoisomerase I can be involved in repression mechanisms [55], this does not occur because it removes negative supercoils, but rather because topoisomerases can block the promoter. In our model, the stochastic behaviour of topoisomerases can also block promoters or elongating RNAPs, but the enzymes eventually unbind, allowing transcription to resume.

## 5 Conclusion

In this work, we introduced TORCphysics, a simulation framework based on physical models that describe DNA–enzyme interactions to investigate supercoiling-driven transcription in genetic circuits. The novelty that TORCphysics offers is versatility, enabling users to define distinct activity models for different types of macromolecules and DNA sites, such as promoters. This flexibility allows the integration and reproduction of diverse regulatory behaviours. TORCphysics also introduces stochastic topoisomerase activity that depend on local superhelical density, effects not previously considered in combination.

Our results highlight the importance of incorporating these key factors, but the model could still be further refined to include more complex mechanics. In particular, our gyrase model tends to overestimate the introduction of twist (e.g., Figure 4b), a discrepancy that might be mitigated by more refined physical models developed in conjunction with experiments. For instance, the current linear model of gyrase activity does not account for ATP hydrolysis, which is essential for DNA supercoiling, and assumes constant ATP availability, an assumption that does not always reflects *in vivo* systems. Incorporating energy-dependent models coupled to experiments where ATP availability is monitored or controlled, could not only help resolve the discrepancies observed, but also enable the investigation of more complex behaviours, such as a gyrase-ATP dependency, ATP/ADP ratio and ATP-independent relaxation of negatively-supercoiled DNA by DNA gyrase.

We also leveraged ChIP-sequencing data from Sutormin et al. [24] to parameterise and simulate the interactions between DNA topoisomerase I and transcribing RNAPs, where we estimated the RNAP twist injection rate into the DNA. However, this parameter should be investigated from experiments specifically designed to probe the interaction of DNA and transcribing RNAPs, as its magnitude may also depend on transcript length and crowding. Additionally, genes involved in membrane-tethered transcription, such as *tetA*, commonly used in pBR322-based plasmids, may exhibit significantly higher twist injection due to constraints on RNAP rotation [56]. These findings highlight the importance of more multimodal modeling approaches, where bioinformatics datasets can serve as a bridge between theoretical models and *in vivo* measurements, while complementary single-molecule experiments are essential for quantifying key DNA-enzyme interactions. Such studies can support the development of models with predictive power, potentially advancing applications in areas like synthetic biology and engineering biology. In this context, we propose TORCphysics as a suitable platform for testing and refining these types of experimental hypotheses.

Lastly, the methodology developed to infer open-complex formation modulation and promoter kinetic properties can also be applied to experimental systems such as those studied by Boulas et al. [18], to further investigate the interplay between transcription and supercoiling. Additionally, current models represent RNAP dynamics using a constant transcription speed of 30 bp/s, which is unlikely in realistic biological conditions, and TORCphysics provides a platform for investigating such questions. TORCphysics demonstrates strong potential as a tool for studying these complex regulatory mechanisms and offers flexibility for expansion. For instance, it can be adapted to incorporate additional models and explore a broader range of transcriptional regulation processes, including canonical repression, DNA looping, and beyond.

## 6 Data availability

The TORCphysics code repository is publicly available on GitHub at: https://github.com/Victor-93/TORCphysics. The dedicated code branch used to generate all results presented in this manuscript, including the processed data and analysis scripts, is available at: https://github.com/Victor-93/TORCphysics/tree/TORCphysics_paper. Further details about the contents of these repositories can be found in Supplemental Material Section 2.

