## Supplemental Material for "TORCphysics: A physical model of DNA-topology-controlled gene expression"

### 1 Overview

Here we present the detailed methodology used in TORCphysics, which includes:

- TORCphysics code repository — see Section 2.
- Parameterising promoter melting energy using the SIST algorithm — see Section 3..
- Marko's elastic model of supercoiled DNA — see Section 4.
- Calculating global superhelical density from experimental kinetic parameters of DNA topoisomerases — see Section 5.
- Elastic function and spacer model equivalency — see Section 6.
- Supplementary Figures - see Section 7.
- Supplementary Tables - see Section 8.

### 2 TORCphysics Repository

TORCphysics is written in Python, and the source code is available on GitHub at `https://github.com/Victor-93/TORCphysics`. The repository includes the source code, documentation, and three Jupyter notebooks examples.

TORCphysics can be installed using the command pip install TORCphysics, or downloaded manually as a ZIP file. The software can be executed via the command line or used in Python scripts.

The repository provides three basic usage examples as Jupyter notebooks, located in the Examples/ directory:

- Example_1.ipynb — Run single gene simulations and analysis.
- Example_2.ipynb — Run multiple simulations with statistical analysis.
- Example_3.ipynb — Define custom enzyme/site models using built-in models.

We also provide a dedicated branch, `TORCphysics_paper`, to reproduce the results presented in this manuscript. This branch is accessible at `https://github.com/Victor-93/TORCphysics/tree/TORCphysics_paper`. The necessary scripts and data for reproducing, processing, and plotting the results are located in the Experiments/ directory. Each subdirectory contains a `README` file with instructions for running the scripts.

These scripts are organised into three directories:

- `Topokinetics/` — Contains scripts for reproducing the stochastic topoisomerase activity experiment.
- `TopoITracksRNAP/` — Contains scripts for the RNAP tracking by Topoisomerase I experiment.
- `Genearchitecture/` — Contains scripts for the gene architecture experiments, including the preprocess of running SIST to parameterise promoter melting energies.

### 3 Parameterising Promoter Melting Energy with SIST

The one-step and three-step superhelical-dependent transcription models introduce the function $U_{\text{melt}}$, which represents the energy required to melt the promoter. This function is used to modulate the binding of RNAPs in the one-step superhelical-dependent transcription model and to modulate the open-complex formation in the three-step transcription model. Here, we describe the methodology used to parameterise the free energy function $U_{\text{melt}}(\sigma, s_p)$ as a function of the superhelical density $\sigma$ and promoter sequence $s_p$ within the genomic sequence $s$. More specifically, we implement the Superhelically Induced Duplex Destabilization (SIDD) model from the SIST algorithm [1] to calculate energy profiles.

Given a genomic sequence $s$, the SIST algorithm can calculate the free energy profiles $G(s_i, \sigma)$ that represent the relative stability of a given base-pair $i$ to transition to strand separation at a given superhelical density $\sigma$. A high value of $G(s_i, \sigma)$ indicates a stable base-pair with low probability of strand separation, while a low value indicates a destabilization and a high probability of melting,

For the gene architecture experiments, we parameterise $U_{\text{melt}}$ for the week, medium and strong promoters, by flanking each promoter $s_p$ by 250 GC base-pairs on each side. This results in the genomic sequence $s = s_{GC} + s_p + s_{GC}$ with $s_{GC}$ representing the 250 flanking GC sequences.

We then run SIST across a range of superhelical densities from $-0.2 \leq \sigma \leq 0.0$ to obtain the free energy profiles $G(s_i, \sigma)$. Figure S2 shows these free energy profiles for the three promoters.

To capture the promoter response to melting, we calculate the average energy $\bar{G}(s_p, \sigma)$ in the promoter region. This average energy typically follows a sigmoidal curve (see Figure S3). We then fit the following sigmoidal function:

$$U_{\text{melt}}(\sigma) = a + \frac{b}{1 + \exp\left(-\frac{\sigma - \sigma_m}{\epsilon_m}\right)} \quad (1)$$

Here, the parameters $a, b, \sigma_m, \epsilon_m$ are sequenced-dependent. The threshold ($\sigma_m$) and width ($\epsilon_m$) are the parameters directly used in the rate equations $k_{\text{on}}$ and $k_{\text{open}}$ from the one-step and three-step superhelical-dependent transcription models, respectively. The fitted ($U_{\text{melt}}$) functions for each promoter are shown in Figure S3. Note that that $U_{\text{melt}}$, as well as parameters $a$ and $b$, are in kcal/mol units, while $\sigma_m$ and $\epsilon_m$ are dimensionless.

Since we use $U_{\text{melt}}$ to modulate rates in the transcription models, we define a simplified, dimensionless form:

$$U'_{\text{melt}}(\sigma) = \frac{\mu}{1 + \exp\left(-\frac{\sigma - \sigma_m(s_p)}{\epsilon_m(s_p)}\right)} \quad (2)$$

Where we have introduced the dimensionless parameter $\mu$ and discarded $a$ and $b$.

Given a rate $k$ (as in the transcription models), it can be modulated through $U'_{\text{melt}}$ by:

$$k(\sigma) = k \exp\left(-U'_{\text{melt}}(\sigma)\right) \quad (3)$$

We choose $\mu$ such that the rate $k$ is reduced to 10% of its maximum activity when $\sigma >> \sigma_m$. In other words:

$$k(\sigma >> \sigma_m) = k \exp(-\mu) = (0.10)k \quad (4)$$

this results in $\mu \approx 2.3$.

This approach can be used not only to model melting response of any promoter, but can also be applied to modeling of strand separation susceptibility of any DNA region. It is important to note, however, that the resulting energy profiles and thus the parameterised $U_{\text{melt}}$ (see Figures S2 and S3, respectively) are highly sensitive to the surrounding DNA sequence context. For instance, sequences with AT-rich regions may be susceptible to melting, potentially competing with strand-separation at the promoter itself, thereby considerably altering both energy profiles and promoter activity [2]. When the sequence context is unknown, we recommend using the flanking GC sequence method as an approximation. Nonetheless, further investigation is needed to fully understand how local sequence context influences DNA energetics and transcriptional regulation.

### 4 Marko's Elastic Model of Supercoiled DNA

In TORCphysics, the velocity of transcribing RNAPs is modeled using a torque-dependent form. To compute torque as a function of superhelical density, TORCphysics employs Marko's elastic model of supercoiled DNA [3]. For a DNA segment held under a constant stretching force $f$ (in pN) and a superhelical density $\sigma$, the torque $\tau$ is given by:

$$\tau = \begin{cases} \dfrac{c_s}{\omega_0'}\sigma, & \text{if } |\sigma| < |\sigma_s| \\ \dfrac{\sqrt{2pg/(1-p/c_s)}}{\omega_0'}, & \text{if } |\sigma_s| < |\sigma| < |\sigma_p| \\ \dfrac{p}{\omega_0'}\sigma, & \text{if } |\sigma| > |\sigma_p| \end{cases} \tag{5}$$

The torque scales linearly with $\sigma$ until $|\sigma| = |\sigma_s|$. In this regime the DNA exists purely in the form of twist. In the coexisting regime $|\sigma_s| < |\sigma| < |\sigma_p|$, where the DNA exists in the form of twist and writhe, the torque remains constant. Beyond $|\sigma| = |\sigma_p|$, supercoiling is stored entirely as writhe (plectonemic form).

The parameter $\omega_0' = \omega_0/.34$ (rad/nm) corresponds to the contour-length rate of rotation, with 0.34 nm being the contour length of a single base-pair in relaxed B-DNA form. The parameter $p = k_B T P \omega_0'^2$ (pN) (with $P$ in length units) describes the twist stiffness of writhed DNA. The free energies per length (pN) for stretching ($g$) and twisting ($c_s$) are given by:

$$g = f - \sqrt{\frac{k_B T f}{A}} \tag{6}$$

$$c_s = c\left(1 - \frac{C}{4A}\sqrt{\frac{k_B T}{A f}}\right) \tag{7}$$

where $A$ (nm) is the bending persistence length, $C$ (nm) the twist persistence length, and $c = k_B T C \omega_0'^2$ (pN) is the twist stiffness of DNA.

The critical values $\sigma_s$ and $\sigma_p$, which define the boundaries of the three supercoiled regimes, are calculated as:

$$|\sigma_s| = \frac{1}{c_s}\sqrt{\frac{2pg}{1 - p/c_s}} \tag{8}$$

$$|\sigma_p| = \frac{1}{p}\sqrt{\frac{2pg}{1 - p/c_s}} \tag{9}$$

This elastic model is valid for relatively low stretching forces (a few pN). In TORCphysics, we assume all stretching forces are low and constant. The specific parameter values used in this model are listed in Table S1.

It is important to note that TORCphysics only considers supercoiling in the form of twist, whereas Marko's model accounts for writhed states. At typical physiological superhelical densities ($\sigma \in [-0.06, -0.04]$), DNA tends to exist in the coexistence regime according to Marko's model with parameters from table S1, where $\sigma_s$ is relatively low and $\sigma_p$ corresponds to highly supercoiled states. We consider this model suitable for TORCphysics as this leads to an approximately constant torque for most scenarios. In cases of hypernegatively supercoiled DNA, RNAPs would stall immediately, due to the sigmoidal form of the RNAP velocity equation.

### 5 Calculating Global Superhelical Density from Experimental Kinetic Parameters of DNA Topoisomerases

In this section, we present the methodology for constructing the reference curves for the change in DNA superhelical density, used to calibrate the models that simulate the stochastic activity of topoisomerases. In general, the process consists integrating reaction curves using the kinetic parameters measured by Wang et al. [4], and then inferring the global superhelical density.

#### 5.1 Kinetic curves integration

Simple enzyme kinetics can be described by the Michaelis-Menten equation:

$$v = v_{\max}\frac{S}{K_M + S} \tag{10}$$

$$k_{\text{cat}} = \frac{v_{\max}}{E} \tag{11}$$

$$v = \frac{dP}{dt} = -\frac{dS}{dt} \tag{12}$$

Here $v$ represents the velocity or rate of product formation $P$, $S$ denotes the substrate concentration, $E$ the enzyme concentration, $P$ the product concentration, $v_{\text{max}}$ the maximum velocity, $K_M$ the Michaelis constant, and $k_{\text{cat}}$ the catalytic rate constant. This equation describes how reaction rates vary with changes in enzyme and substrate concentration. The general reaction scheme of an enzyme-catalyzed reaction is illustrated in figure S8a).

Given an initial substrate concentration and a set of kinetic parameters ($v_{\text{max}}, K_M, k_{\text{cat}}$), we can numerically integrate the Michaelis-Menten equation using the Euler method:

$$P_{i+1} = P_i + v_i \Delta t \tag{13}$$

$$S_{i+1} = S_i - v_i \Delta t \tag{14}$$

By iterating and updating these simple set of equations, we can generate the resulting kinetic curves.

#### 5.2 Topoisomerase I (topo I) kinetics

The reaction scheme for the relaxation reaction catalyzed by *E. coli* DNA topoisomerase I (topo I), as proposed in [4], is shown in Figure S8b. According to this scheme and following the Michaelis-Menten equation10, the substrate would be the concentration of supercoiled DNA, to which topo I binds, resulting in the production of relaxed DNA. In their kinetic study, an initial plasmid concentration of 0.75nM supercoiled DNA and a constant topo I concentration of 17.0 nM were utilized. By incorporating these concentrations and the kinetic parameters derived in their study (see Table S2), we were able to numerically integrate and obtain the relaxation curves shown in figure S9, where topo I relaxes supercoiled DNA, transitioning the total plasmid concentration from supercoiled DNA to relaxed DNA.

We can associate the reaction curves to the change in superhelical density in the plasmid by assuming a linear relationship between the concentration of relaxed DNA and the superhelical density:

$$\sigma(t) = a + R(t)b \tag{15}$$

where $R$ is the concentration of relaxed DNA (product) as a function of time. When the superhelical density is zero, the concentration of relaxed DNA corresponds to the total plasmid concentration of .75nM. We then assume that at the superhelical density of -0.11 suggested as the maximum superhelical density induced by DNA gyrase according to previous observations [5], the total plasmid concentration shifts to supercoiled DNA, hence $R = 0$. Any state in between corresponds to a mixture of superhelical and relaxed DNA. Given these conditions, the previous relationship takes the form:

$$\sigma(t) = -0.11 + 0.11 \frac{R(t)}{0.75\text{nM}} \tag{16}$$

Figure S9 shows the resulting relaxation curve in terms of the concentration of relaxed DNA (panel a), the concentration of supercoiled DNA (panel b), and the corresponding superhelical density calculated using equation 5.2 for the where topo I interacts solely with supercoiled DNA (red curve). For this case, we set the initial superhelical density to $\sigma = -0.11$, which corresponds to a fully supercoiled DNA state ($R = 0$).

#### 5.3 Gyrase kinetics

Similar to topo I, the reaction scheme for the supercoiling-induced reaction catalyzed by *E. coli* DNA Gyrase is shown in figure S8c, as proposed by Wang et al. [4]. According to this kinetic pathway, gyrase binds the relaxed DNA substrate, producing supercoiled DNA, and releasing the gyrase enzyme. Although *E. coli* gyrase has two substrates, relaxed DNA and ATP, we do not consider the gyrase-ATP complex formation in this study. Instead, we assume that gyrase alone interacts with relaxed DNA. However, it is important to note that the kinetic parameters obtained in [4] and utilized in this study (see table S2, maintained the ATP concentration at 1.75 mM during their assay.

By implementing and integrating the Michaelis-Menten equation (see equations 10-12, and utilizing the gyrase kinetic parameters obtained in [4] (see table S2), we are able to generate the supercoiled-induced curves for DNA Gyrase interacting with relaxed DNA (see figure S9b, represented by the blue curve). We start from a relaxed DNA state corresponding to $R = 0.75$nM and $\sigma = 0$. By applying the relationship between the superhelical density and the concentration of relaxed DNA, we can infer the superhelical density using equation 5.2. Figure S9c (blue curve) illustrates the resulting superhelical density for DNA gyrase interacting with relaxed DNA.

#### 5.4 Both topo I and gyrase kinetic curve

In addition to the scenarios involving DNA gyrase or topo I interacting independently with DNA, we simulate two cases where both enzymes interact simultaneously with the DNA by integrating the Michaelis-Menten equation 10. Here, topo I acts on the supercoiled DNA substrate ($C$)) and produces relaxed DNA ($R$), while gyrase acts on relaxed DNA substrate and produces supercoiled DNA (see figure S8b-c). The corresponding set of kinetic equations is as follows:

$$C_{i+1} = C_i + (v_{\text{gyrase}} - v_{\text{topoI}})\Delta t \tag{17}$$

$$R_{i+1} = R_i - (v_{\text{gyrase}} - v_{\text{topoI}})\Delta t \tag{18}$$

The rates $v_{\text{gyrase}}$ and $v_{\text{topoI}}$ are determined by substituing the corresponding gyrase and topo I kinetic parameters and substrates (relaxed DNA and supercoiled DNA) into equation 10. We initiate the kinetic simulations at the superhelical level of 0, corresponding to a relaxed state with concentration of relaxed DNA of 0.75nM and 0 for supercoiled

DNA, and at the superhelical level of $-0.11$, corresponding to a concentration of supercoiled DNA of 0.75nM and 0 for relaxed DNA (refer to figure S9a-b). Figure S9c displays the corresponding superhelical density for both cases (green and purple curves), where both conditions tend to the plateau with a value of approximately -0.046, approximating physiological values of in vivo superhelical density in *E. coli* [6].

### 6 Elastic function and spacer model equivalency

The three-step superhelical-dependant transcription model used in TORCphysics incorporates an elastic function $G_{\text{elastic}}$ to emulate the closed-complex formation. A previous model proposed by Forquet et al. [7] predicts the relative activation of promoters based on their spacer length and orientation as a function of DNA supercoiling. Here, we demonstrate that both models are equivalent.

The spacer length model is defined as:

$$G_{\text{spacer}}(\sigma, n) = \frac{n}{2} k_\theta \left( \frac{\theta_p}{n} - \alpha_0 (1 + \sigma) \right)^2 \tag{19}$$

where $n$ is the spacer length, $k_\theta = 71.4\,\text{kBrad}^{-2}$ is the DNA sequence twist stiffness, $\alpha_0 = 34$ is the average twist angle, and $\theta_p$ is the optimal twist angle between the -35 and -10 promoter regions.

The optimal twist angle, $\theta_p$, can be related to the optimal superhelical density $\sigma_0 = -0.06$ as follows:

$$\theta_p = n\alpha_0(1 + \sigma_0) \tag{20}$$

Thus, the spacer length model becomes:

$$G_{\text{spacer}}(\sigma, n) = n k_\theta \alpha_0^2 \frac{(\sigma - \sigma_0)^2}{2} \tag{21}$$

We associate the TORCphysics elastic function with the spacer length model as follows:

$$G_{\text{elastic}}(\sigma, \sigma_e, \epsilon_e) = G_{\text{spacer}}(\sigma, n) \tag{22}$$

$$\frac{(\sigma - \sigma_e)^2}{2\epsilon_e^2} = n k_\theta \alpha_0^2 \frac{(\sigma - \sigma_0)^2}{2} \tag{23}$$

With this, the elastic model takes the form of the spacer length model with $\sigma_e = \sigma_0 = -0.06$ and $\epsilon_e = \frac{1}{\sqrt{n k_\theta \alpha_0^2}}$.

### 7 Supplementary Figures

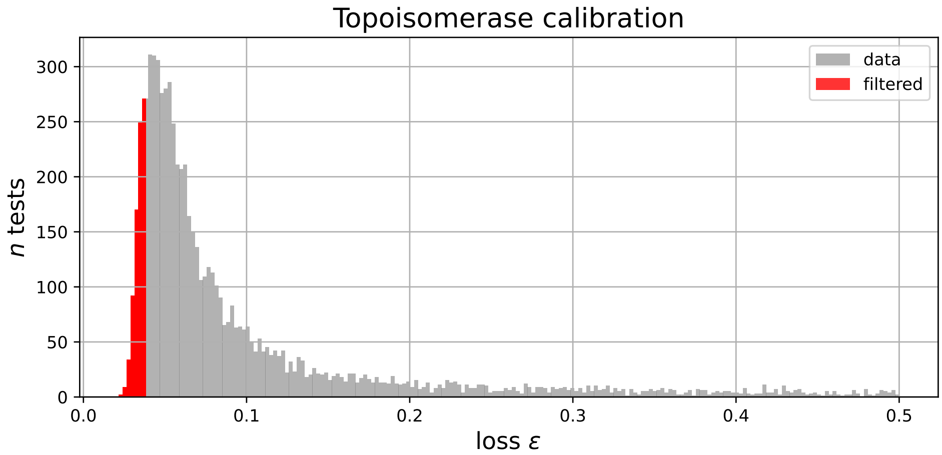


Figure S1: Distribution of losses $\epsilon$ resulting from the calibration of the stochastic activity of topoisomerases. Losses are calculated according the objective function used to evaluate the model. Red bins indicate the top 5% of best parameter sets out of 8,000 random tests.

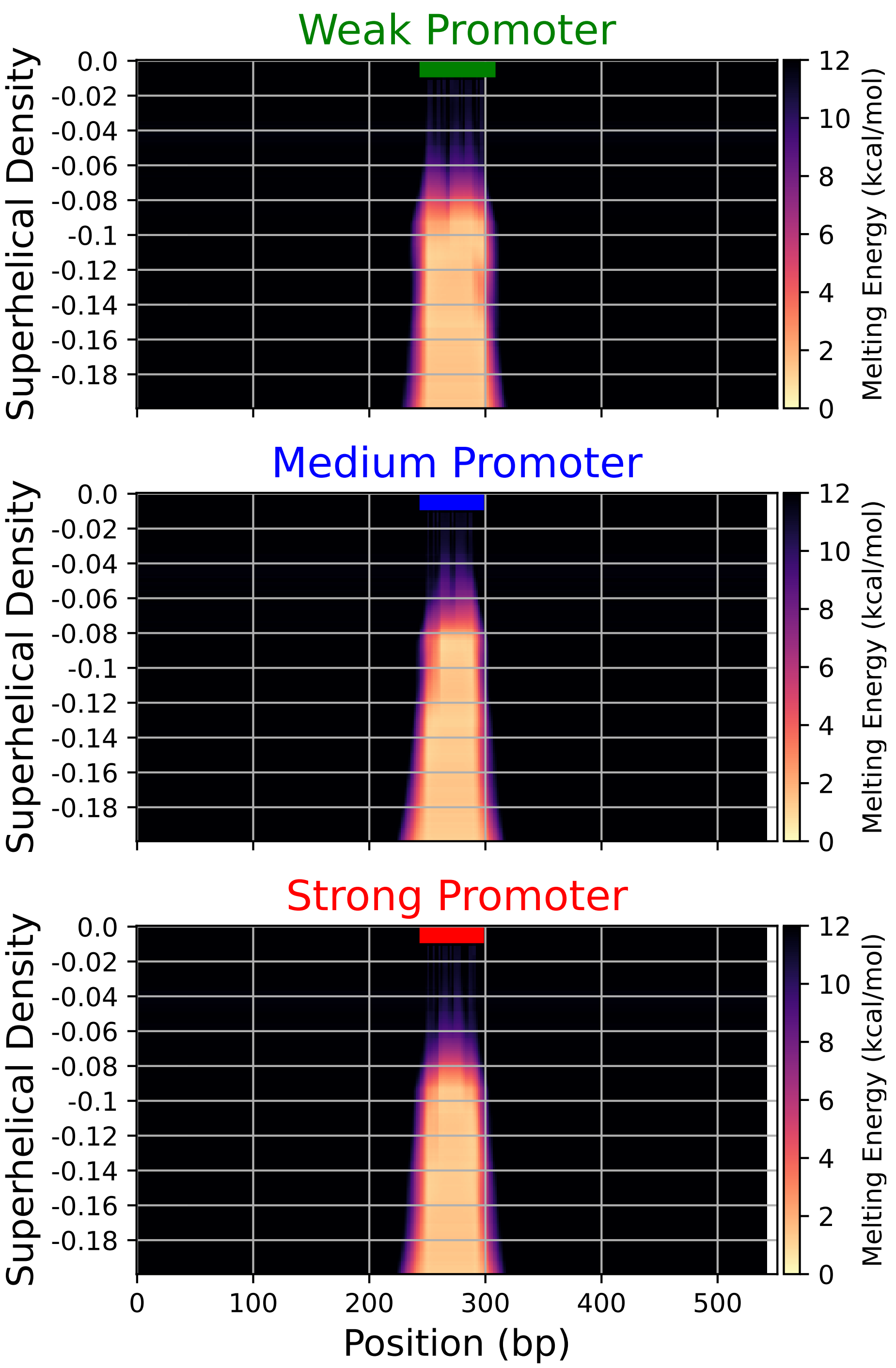


Figure S2: Melting energy profiles ($G(s, \sigma)$) for the weak, medium and strong promoters flanked by 250 GC sequences on each side. The profiles were calculated using SIST. The colored lines at the top of each subplot indicate the promoter positions: green for the weak promoter, blue for the medium promoter, and red for the strong promoter.

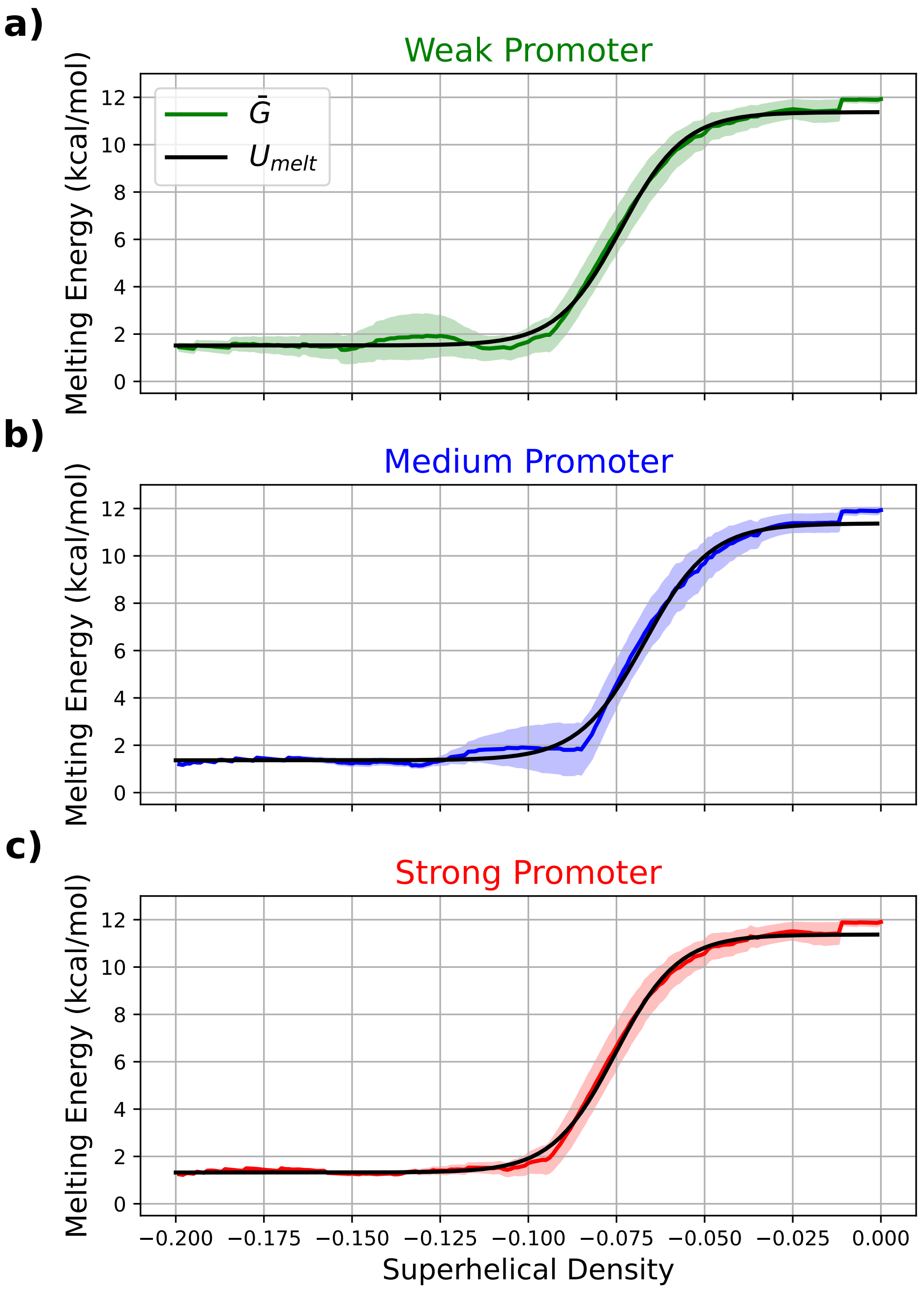


Figure S3: Promoter response $U_{\text{melt}}$ obtained by fitting equation 1 to the averaged melting energy $\bar{G}$, for the weak (green), medium (blue) and strong (red) promoters. The promoter sequences were flanked by 250 GC base-pairs on each side for this case. However, in realistic scenarios, the response depends on sequence context. Energies are averaged around the -10 region, where small standard deviations indicate that the energy required for strand separation do not vary greatly in these regions.

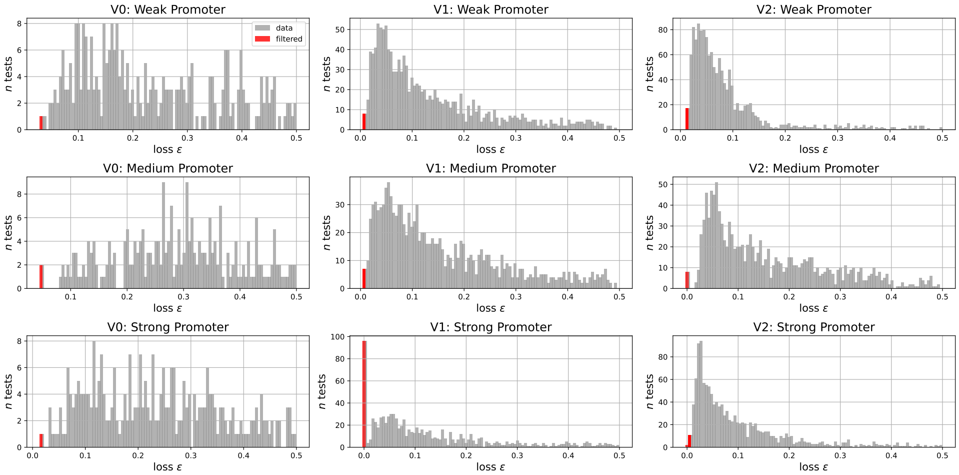


Figure S4: Distribution of losses resulted from the gene architecture calibration process for the three models V0, V1, and V2, across the weak, medium, and strong promoters. The best parameter set is within the red bin. A total of 300 random tests were run for V0, and 1,300 for both V1 and V2.

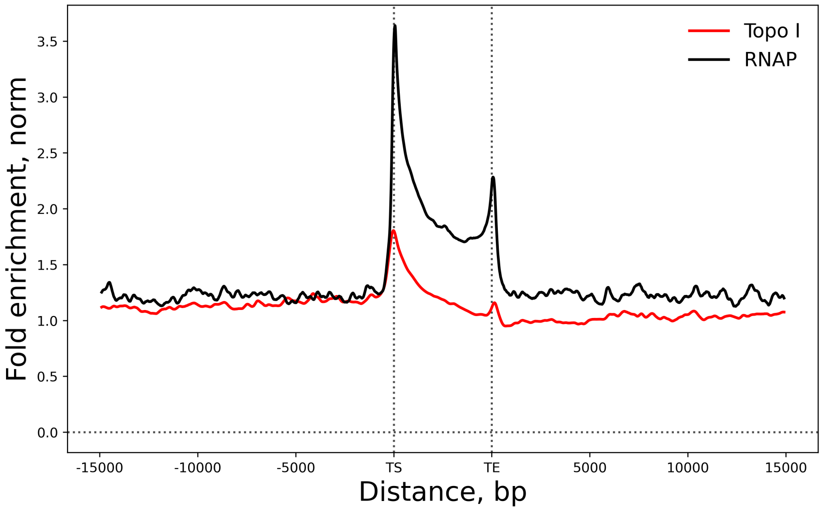


Figure S5: Topoisomerase I fold enrichment and normalized RNAP signal, obtained from Chip-Seq data used in Sutormin et al. [8]. Rather than directly using this data in TORCphysics, the model aims to reproduce equivalent averaged fold enrichment of topoisomeorase I within the TU ($\mathcal{F}_{\text{exp}} = 1.24$), the correlation coefficient between the both curves ($\rho_{\text{exp}} = 0.94$). Transcription unit is defined as the region between the transcription start site (TS) and transcription termination site (TE). TORCphysics simulates an equivalent system composed by a topological domain with a single gene (TU) at the middle.

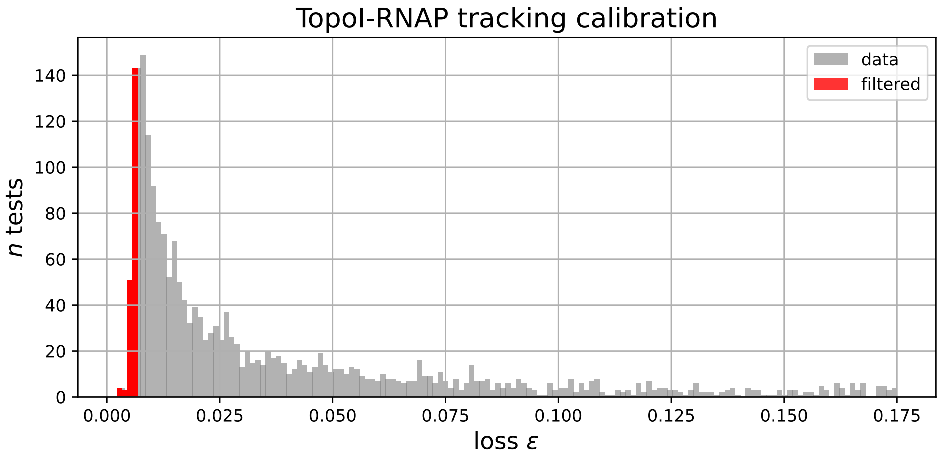


Figure S6: Loss distribution from the calibration of RNAP tracking by topoisomerase I. Loss values were computed using the objective function designed to evaluate how well the model mimics experimental behavior. The top 5% of best parameter sets, out of 3,000 random trials, are highlighted in red.

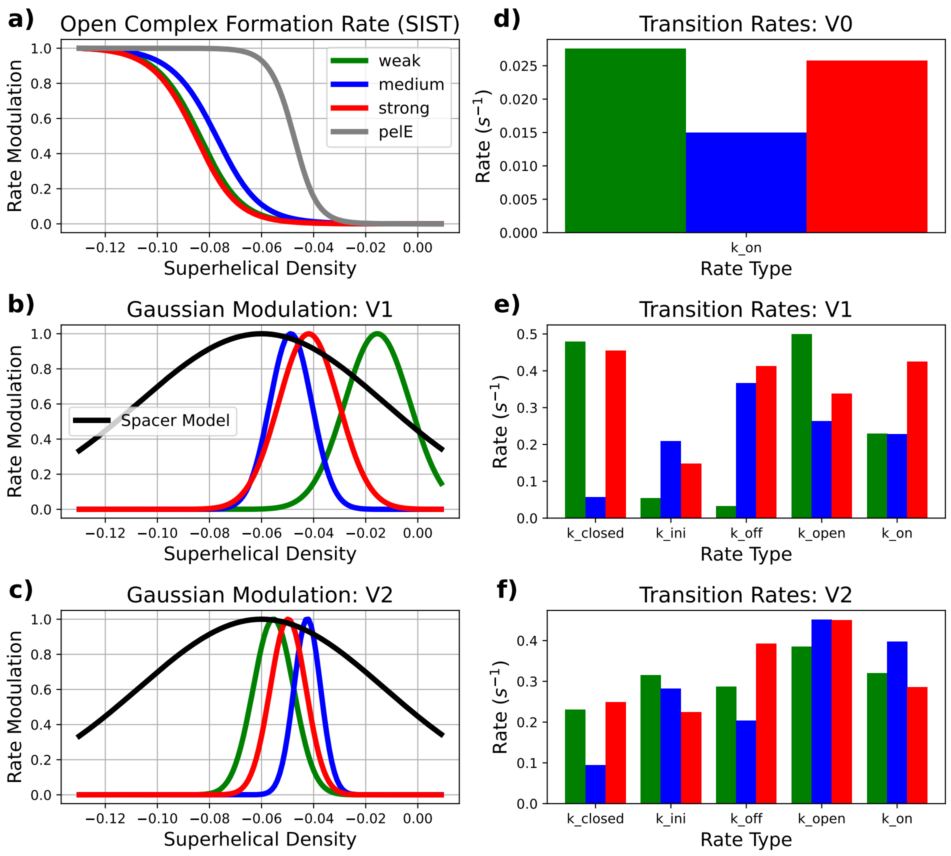


Figure S7: a) Open-complex formation rate modulation for the pelE promoter, calibrated in El Houdaigui et al. [9], alongside the modulation profiles of weak, medium, and strong promoters calibrated using the SIST algorithm [10] and sequences obtained from Boulas et al. [11]. b) and c) Closed-complex formation modulation for the weak, medium, and strong promoters in models V1 and V2, respectively, with the spacer length model from Forquet et al. [7] included for comparison. d) Binding rates $k_{\text{on}}$ for model V0, promoter kinetic rates for models V1 e) and V2 f), resulted from the gene architecture calibration process.

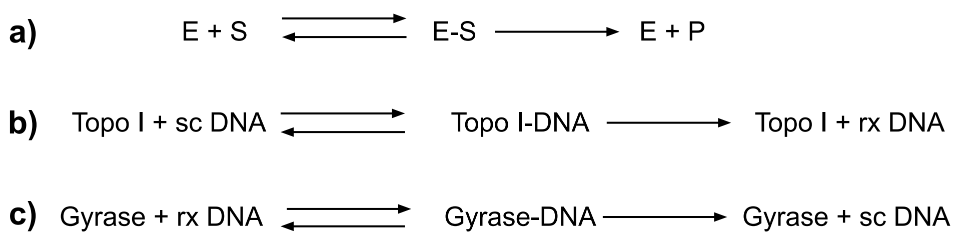


Figure S8: Topoisomerases kinetic reaction scheme in *E. coli* as proposed by Wang et. al. [4]. a) General kinetic pathway described by the Michaelis-Menten equation. Enzyme E binds substrate S to form the complex E-S, followed by product P production and enzyme E release. b) Kinetic pathway of topoisomerase I acting on supercoiled DNA. Topoisomerase I (topo I) binds supercoiled DNA substrate (sc DNA) to form DNA-topoI complex, resulting in relaxed DNA production and topo I release. c) Reaction scheme of DNA gyrase acting on relaxed DNA. Gyrase binds relaxed DNA to form DNA-Gyrase complex, leading to supercoiled DNA production. ATP concentration remains constant in this reaction scheme.

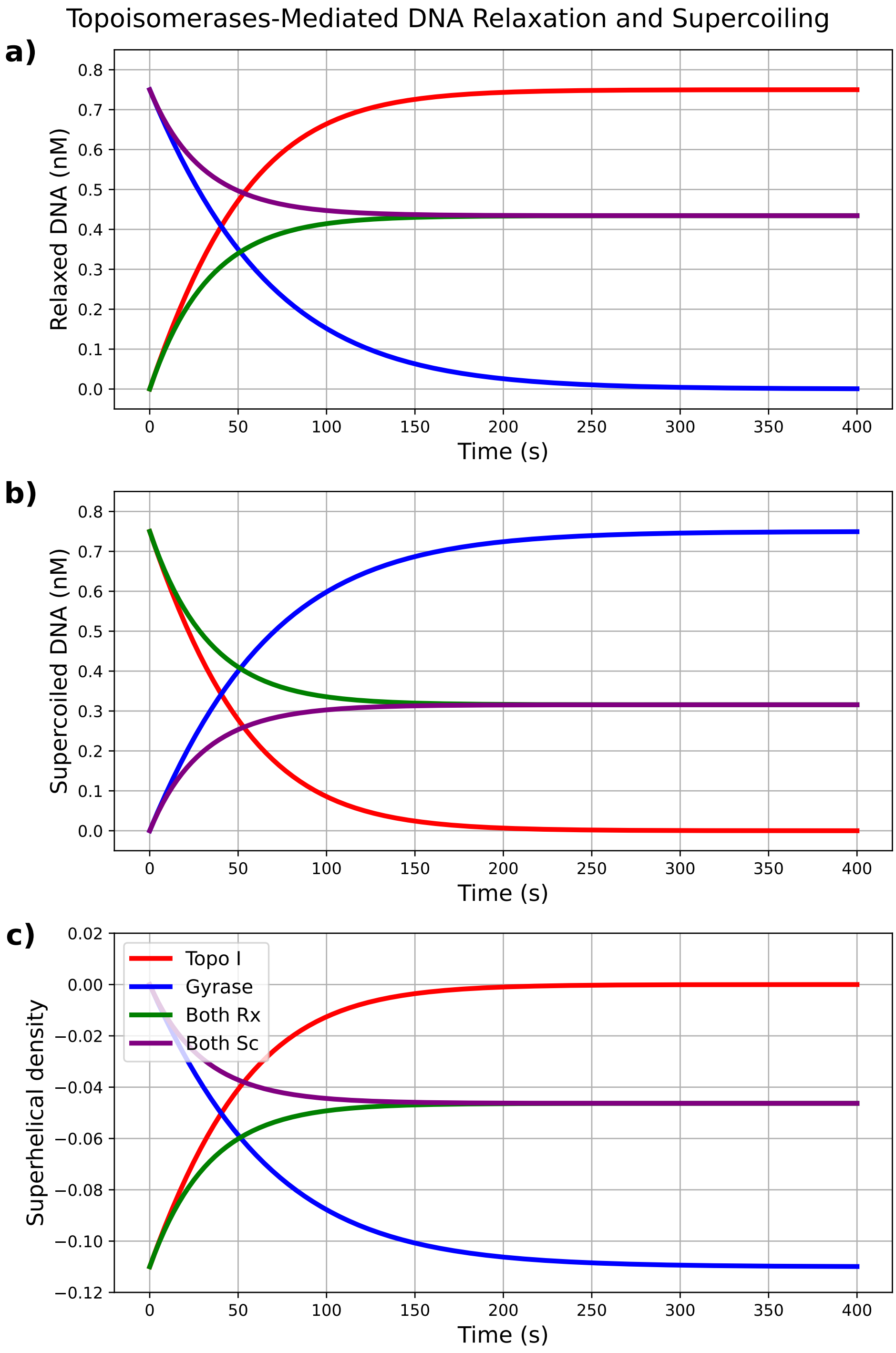


Figure S9: Time courses of relaxation (a) and supercoiling (b) reactions catalyzed by *E. coli* DNA topoisomerase I (topo I) in red, DNA gyrase in blue, both enzymes acting simultaneously on DNA starting from a supercoiled state in green, and starting from a relaxed state in purple, obtained through the integration of the Michaelis-Menten equation. Panel (c) illustrates the corresponding superhelical density determined by a linear relationship with relaxed DNA, as shown in equation 5.2, for the three scenarios. The kinetic parameters utilized are taken from [4] and shown in table S2.

### 8 Supplementary Tables

Table S1: Values of parameters used in TORCphysics for calculating torques according to Marko's elastic model of supercoiled DNA [3]. The energies $g$ and $c_s$ are expressed in terms of free energy per unit length (i.e., units of force). The parameters $p$, $c$, $g$, $c_s$, $\sigma_s$, and $\sigma_p$ were computed using the equations provided in Section 4, based on DNA stiffness and structural parameters, as well as temperature.

| **Parameter** | **Value** | **Description** |
|---|---|---|
| $f$ **(pN)** | 1.00 | Stretching forces |
| $\omega_0'$ **(rad/nm)** | 1.76 | Contour-length rate of rotation |
| $T$ **(K)** | 300 | Temperature |
| $A$ **(nm)** | 50 | Bending persistence length |
| $P$ **(nm)** | 24.0 | Twist persisetence length of plectonomic DNA |
| $p$ **(pN)** | 304 | Twist stiffness of plectonomic DNA |
| $C$ **(nm)** | 95.0 | Twist persistence length |
| $c$ **(pN)** | 1206 | Twist stiffness |
| $g$ **(pN)** | .714 | Free energy of stretched DNA |
| $c_s$ **(pN)** | 1042 | Free energy of twisted DNA |
| $\sigma_s$ **(nm)** | 0.0237 | Twist threshold |
| $\sigma_p$ **(nm)** | 0.0813 | Writhe threshold |

Table S2: Steady-State Kinetic Parameters of *E. coli* DNA Topoisomerase I (topo I) and DNA Gyrase, taken from the kinetic study conducted by Wang et al. (2019) [4].

| Parameter | Topoisomerase I | Gyrase |
|---|---|---|
| $k_{\text{cat}}$ ($s^{-1}$) | 2.3 x $10^{-3}$ | 1.1 x $10^{-3}$ |
| $K_M$ (nM) | 1.5 | 2.7 |
| $v_{\max}$ (pM/s) | 40 | 50 |
| $E$ (nM) | 17 | 45 |

Table S3: Promoter sequences used in this study for the gene architecture sections, taken from Boulas et al. [11]. The -10 promoter regions are highlighted in bold letters.

| Name | Sequence |
|---|---|
| Weak | AAAAAGAGTATTGACTTCGCATCTTTTTGTACC**TATAAT**GTGTGGATAGCGG |
| Medium | TTGACATCAGGAAAATTTTTCTG**CATAAT**TATTTCATATCAC |
| Strong | TTGACATCGCATCTTTTTGTACC**TATAAT**GTGTGGATAGAGT |
